# Impact of IL-21 on Natural Killer cell proliferation and function – a mathematical and functional assessment

**DOI:** 10.1101/2024.01.26.577405

**Authors:** Rosalba Biondo, Indrani Nayak, Nina Möker, Congcong Zhang, William C. Stewart, Salim Khakoo, Jayajit Das

**Affiliations:** School of Clinical and Experimental Sciences, University of Southampton, Southampton, United Kingdom; Steve and Cindy Rasmussen Institute for Genomic Medicine, Abigail Wexner Research Institute, Nationwide Children’s Hospital, Columbus, Ohio; Miltenyi Biotec B.V. & Co. KG, Bergisch Gladbach, Germany; GIG Consulting, Inc, Columbus, Ohio; Biomedical Sciences Graduate Program, The Ohio State University, Columbus, Ohio; Department of Pediatrics, The Ohio State University, Columbus, Ohio; Pelotonia Institute for Immuno-Oncology, The Ohio State University, Columbus, Ohio; The Biophysics Graduate Program, The Ohio State University, Columbus, Ohio

**Keywords:** NK cells, Cytokines, STAT, proliferation, cytotoxicity, linear regression, prediction

## Abstract

Natural killer (NK) cells are currently in use as immunotherapeutic agents for cancer. Many different cytokines are used to generate NK cells including IL-2, IL-12, IL-15 and IL-18 in solution and membrane bound IL-21. These cytokines drive NK cell activation through the integration of STAT and NF-κB pathways, which overlap and synergize, making it challenging to predict optimal cytokine combinations. We integrated functional assays for NK cells cultured in a variety of cytokine combinations with feature selection and mechanistic regression models. Our regression model successfully predicts NK cell proliferation for different cytokine combinations and indicates synergy between STAT3 and NF-κB transcription factors. Use of IL-21 in solution in the priming, but not post-priming phase of NK cell culture resulted in optimal NK cell proliferation, without compromising cytotoxicity or IFN-γ secretion against hepatocellular carcinoma cell lines. Our work provides a mathematical framework for interrogating NK cell activation for cancer immunotherapy.

## Introduction

Natural killer (NK) cells are being increasingly used as an adoptive immunotherapy against tumors. One key challenge in NK cell immunotherapy is the ability to generate sufficient numbers of activated NK cells to be an effective therapeutic. Expansion of NK cells in vitro using specific combinations of cytokines is a promising approach to generate an adequate quality and quantity of NK cells (1–5). NK cells express multiple cytokine receptors for binding to type 1 (IL-2/4/12/15/21), type 2 (such as IFN-γ) and IL-1 (such as IL-18) family cytokines (6). Thus, there are many potential combinations of cytokines that could be used to expand NK cells. A widely used protocol to induce proliferation and maturation of NK cells in vitro is to pre-stimulate NK cells with a combination of cytokines for a short duration, up to 16 hours (“priming”) followed by culture over days where fewer cytokines are used (1). Ex vivo expansion using IL-12/15/18 priming to generate long lived memory NK cells is a promising area of research with clinical benefit obtained in early phase studies (1,7,8). IL-21 is a pleiotropic cytokine that is involved in activation of NK and CD8 T cells and is specifically associated with tissue resident NK cells(9). IL-21 has been shown to induce excellent expansion of NK cells ex-vivo using feeder cell lines expressing membrane bound IL-21, which appears to mitigate the senescence associated with other cytokine regimens (10). However, the use of feeder cell lines has additional regulatory issues relating to the provenance of the cell lines used (11). Combining IL-21 with pre-existing regimes using soluble cytokines provides an alternative strategy to NK cell expansion.

Selecting the correct combinations of cytokines to obtain a NK cell population with a desired response such as large population size or/and greater cytotoxicity against specific target cells is a key challenge. Whilst the effects of single cytokines on NK cells is well understood, the difficulty in choosing regimens using multiple cytokines arises due to our rudimentary understanding of how the different combinations of cytokines combine to stimulate NK cells and induce genes that regulate NK cell proliferation, expression of specific NK cell receptors, and NK cell maturation. NK cells are activated by stimulation of cell surface and cytokine receptors in a combinatorial manner. Signals from different combinations of receptors can synergize to enhance NK cell proliferation and activation.

Stimulation by the type 1 and 2 family of cytokines leads to the phosphorylation of seven signal transducer and activator of transcription (STAT) proteins and stimulation by IL-1 family of cytokines activates nuclear factor-kappa B (NF-κB) transcription factor. These proteins translocate to the nucleus and induce specific gene expressions by binding to promoters/enhancers motifs in the DNA (12). However, there is a substantial overlap between the STAT proteins that are phosphorylated due to different cytokine stimulation, for example, STAT1, STAT3, and STAT4 can be phosphorylated by either IL-2 or IL-12 stimulation (6,12,13). In addition, the STAT proteins can synergize or antagonize as they induce transcription (6). Furthermore, NK cells obtained from different donors can respond differently to the same combination of cytokines. Therefore, relating a combination of cytokines to a specific NK cell response such as proliferation across donors can be challenging. While mechanistic computational models based on ordinary differential equation (ODE) have been developed to describe NK-, T-, and B-cell proliferation under the influence of a variety of cytokines including IL-2 and/or IL-15 (14–18), it is challenging to extend such models to include priming and post-priming by a cocktail of cytokines.

Machine learning-based or mechanistic ODE, as well as stochastic simulation-based models have been explored in relating cytokines with predictive outcomes of pathogenesis, or T cell activation (19–23) and kinetics of phosphorylated STAT proteins and gene expression responses in bone marrow derived macrophages (24) and dendritic cells (25) treated with cytokines over several hours to few days. However, to date, the roles of synergy/antagonism between multiple cytokines in regulating NK cell proliferation remains unexplored. By including the effect of synergy/antagonism between STAT and NF-κB transcription factors induced by multiple cytokines, we address this challenge computationally combined with experiments by developing a predictive framework combining longitudinal measurement of NK cell populations, regression model with LASSO regularization (26), and a filter-style feature selection algorithm (RRelief) (27), commonly used for guiding machine learning methods, that can predict fold expansions in NK cell population using different cytokine combinations.

## Results

### IL-21 induces early rapid proliferation of NK cells

To investigate the effects of different cytokines on proliferation of NK cells, NK cells were purified from 17 donors and treated with 12 different combinations of IL-2, IL-12, IL-15, IL-18, and IL-21 (**Fig. 1A** and **B**). Ten donors (Donor 1 – Donor 10) were treated with cytokine conditions 1-6, four donors (Donor 11 – Donor 14) were treated with cytokine conditions 7-8, and three donors (Donor 15 – Donor 17) were treated with conditions 9-12 (**Table S1A**). The experimental design involved a priming step of 16 hours, followed by a post-priming consisting of a different cytokine combination for up to three days (denoted as post-priming I or PP-I), and finally a maintenance regimen of IL-2 alone or IL-2 in combination with IL-15 (denoted as post-priming II or PP-II), as shown in **Fig. 1B**. Comparisons were made within donor groups only. Cells were cultured for up to 16 days and the average NK cell fold expansion was measured across donors for each cytokine cocktail condition at day 2, 4, 7, 9, 12, 14 and 16 (**Fig. 1C** and **Table S2B**). Direct comparison of condition 2 (IL-12/15/18 priming) and condition 3 (IL-12/15/18/21 priming) showed that positive effects of IL-21 could be seen as early as day four of culture (**Fig. 1C**). By day 9 under all conditions, NK cells had proliferated between 10-15-fold, with the IL-21 priming regimens, condition 3 giving better early proliferation as compared to condition 2 (p<0.05). The effects of IL-21 in general seemed positive by this stage but were less evident in the later stages of culture. However, there was a substantial donor heterogeneity within these responses with one donor having a ∼70-fold proliferation with condition 3, which was approximately 3-fold any other donor. (**Table S1A**).

**Figure 1.**
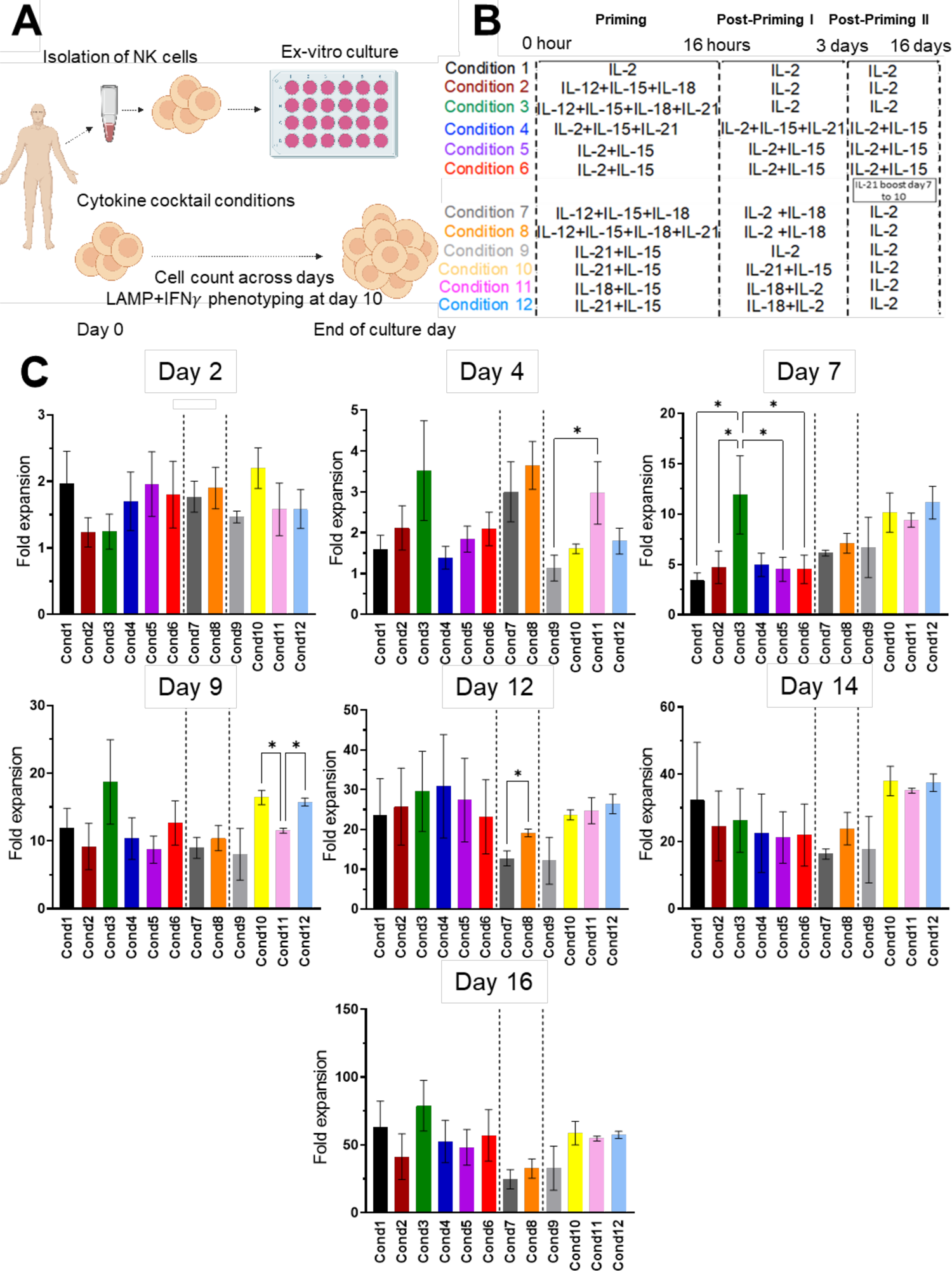
Testing different cytokine conditions for NK cell proliferation. **(A)** NK cells were isolated from the peripheral blood of 17 healthy donors and cultured with different combinations of cytokines with fresh media replenished every 2-3 days. **(B)** Table illustrating the different combinations of cytokines used to culture NK cells. **(C)** Proliferation of NK cells treated with the regimens as illustrated in **B**. Ten donors were treated with conditions 1-6, 4 donors with conditions 7 and 8, and 3 donors with conditions 9-12. Analyses were done only between conditions evaluating the same donors. Data shown as mean ± SEM (*P≤0.05). Data analysed by one-way ANOVA, and Tukey’s multiple comparison test for conditions 1-6 and 9-12, and paired t-test for condition 7 and 8.

To investigate the level of proliferation further, we selected the overall two best IL-21 priming regimes (conditions 3 and 12) and used CFSE to determine the rate of cell division as compared to a control conditions IL-2 alone or IL-12/15/18 priming (**Fig. 2A** and **2B**). Consistent with our initial observations, the difference in proliferation was most clearly seen at day 7 (gray bars) for IL-21 containing regimes (conditions 3 and 12), compared to the control conditions. A higher proportion of NK cells had undergone seven cell divisions by day 7 compared to day 9, for both IL-21 containing condition (population 7 (gray bars) in **Fig. 2B**). The percentage of CFSE-positive NK cells from population 7, decreased from a mean of 40% at day 7 to 25% at day 9 in condition 3, and from 45% at day 7 to 10% in condition 12 (**Fig. 2B**). The greatest loss of NK cells was found in condition 12, which had IL-21 and IL-15 in the priming step, as opposed to IL-12/15/18 plus IL-21, that is inclusion of IL-21 with the full set of cytokines that are in clinical use to induce memory NK cells, which is currently in clinical use (population 7, **Fig. 2B**).

**Figure 2.**
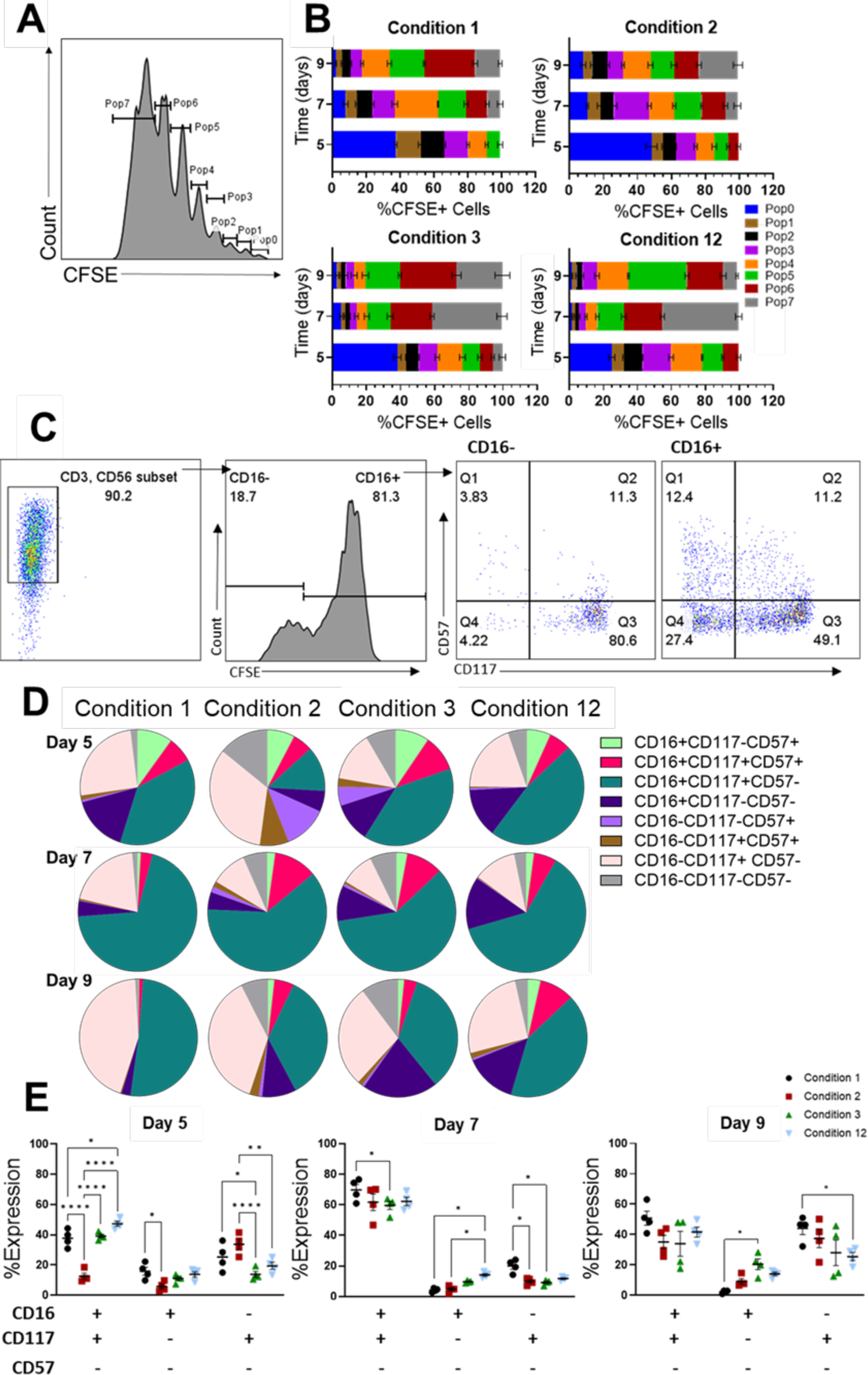
Proliferation and maturation of NK cell subpopulations. NK cells from 4 donors were stained with CellTrace CFSE, cultured with condition 1, condition 2, condition 3, condition 12 and assessed by flowcytometry at day 5, 7 and 9 of in-vitro culture. **(A)** Representative example of gating strategy. NK cells were gated on each cell division, according to the levels of fluorescence of CFSE and this was used to determine the size of each population, plotted in **B**. **(B)** Proliferation of cells, based on their CFSE percentage. The width of each color segment reflects the proportion of the individual populations in the entire CFSE+ cell population. Each division is marked by a difference in CFSE retention by the cells, marking the different populations (pop0 to pop7) within the whole heterogenous cell pool. Data are shown as mean± SEM (N=4). **(C)** Representative example of gating strategy. NK cells population was analysed for the expression of CD16, CD57 and CD117, by flowcytometry. Both CD16- and CD16+ subpopulations were then gated according to CD57 and CD117 expressions. (**D**) Proportions of NK subpopulations according to the expression of CD16, CD117, and CD57, at day 5, 7 and 9. (**E**) Comparison of the most abundant NK subpopulations at day 5, 7 and 9. Data shown as mean± SEM (n=4), analysed by two-way ANOVA with Tukey’s multiple comparison test (*P≤0.05, **P≤0.001, ****P≤0.0001). (**D**-**E**) Values are expressed as a percentage of either CD16- or CD16+.

The proliferating NK cells subpopulations, were analyzed for the expression of the maturation markers CD117 (immature), CD16 (mature), and CD57 (terminally differentiated) (**Fig. 2D** and **E**). During the early stages of the culture, (day 5, **Fig. 2D**) NK cells maturation is heterogeneous, with CD16+CD117+CD57-(dark green in **Fig. 2D**), CD16+CD117-CD57-(purple in **Fig. 2D**) and CD16-CD117+CD57+ (pink in **Fig. 2D**) being the most prominent phenotypes. These subpopulations were affected differently by the different culture conditions throughout the *in vitro* culture (**Fig. 2D**). By day 7, over 60% of the NK cell population shows a phenotype which is predominantly CD16+CD117+CD57-(dark green in **Fig. 2D**). A closer comparison of the conditions at day 7, shows that condition 1 yields the highest proportion of CD16+CD117+CD57-NK cells, accounting for approximately 70% of the population. Condition 2, condition 3, and condition 12 demonstrated comparable results, exhibiting approximately 60% CD16+CD117+CD57-NK cells. (**Fig. 2E**). Over the course of the culture period (at day 9), we observed the emergence of CD16-CD117+CD57-(pink in **Fig. 2D** and **E**) population in conditions 1 and 2 (no IL-21 in the priming phase), comprising approximately 40% and 30% of the population, respectively (**Fig. 2E**). Conversely, the IL-21 containing cultures (condition 3 and condition 12) exhibited a higher proportion of CD16+CD57-CD117+/- NK cells, representing over 55% of the whole population (green and purple in **Fig. 2D** and **Fig. 2E**). Thus IL-21 supports the generation of a key sub-population of cytotoxic CD16+ NK cells.

### IL21-STAT3 signaling determines proliferation

To investigate the role of JAK/STAT and NF-κB signaling pathways in IL-21 mediated NK cell proliferation, NK cells were cultured using condition 2 (IL-21 absent) or condition 3 (IL-21 present) in the presence of either of STAT3 (S31-201) or NF-κB (PS1145) inhibitors and proliferation was measured. These results show that proliferation was dependent on both pathways for both conditions (**Fig. 3A** and **B**). However, in condition 2, the effect of STAT3 inhibition was greater than that for NF-κB (p=0.0002), whereas the reverse was true for condition 3 (p=0.0127) (**Fig. 3A**). Direct comparison of condition 2 and 3 on individual donors confirmed that inhibition of the NF-κB pathway had a more profound effect on condition 3 than on condition 2 (p<0.0001), whereas the reverse was true for STAT3 (p=0.0088) (**Fig. 3B**). Phosphorylation of STAT3 in the absence of any inhibitors was found to be more evident in condition 3 than condition 2 (**Fig. 3C**). Overall, this suggests that IL-21 exerts its additional pro-proliferative effect predominantly via STAT3. However, due to the complexity of cytokine signaling in NK cells we took a modelling approach to further interrogate how they may affect NK cell proliferation.

**Figure 3.**
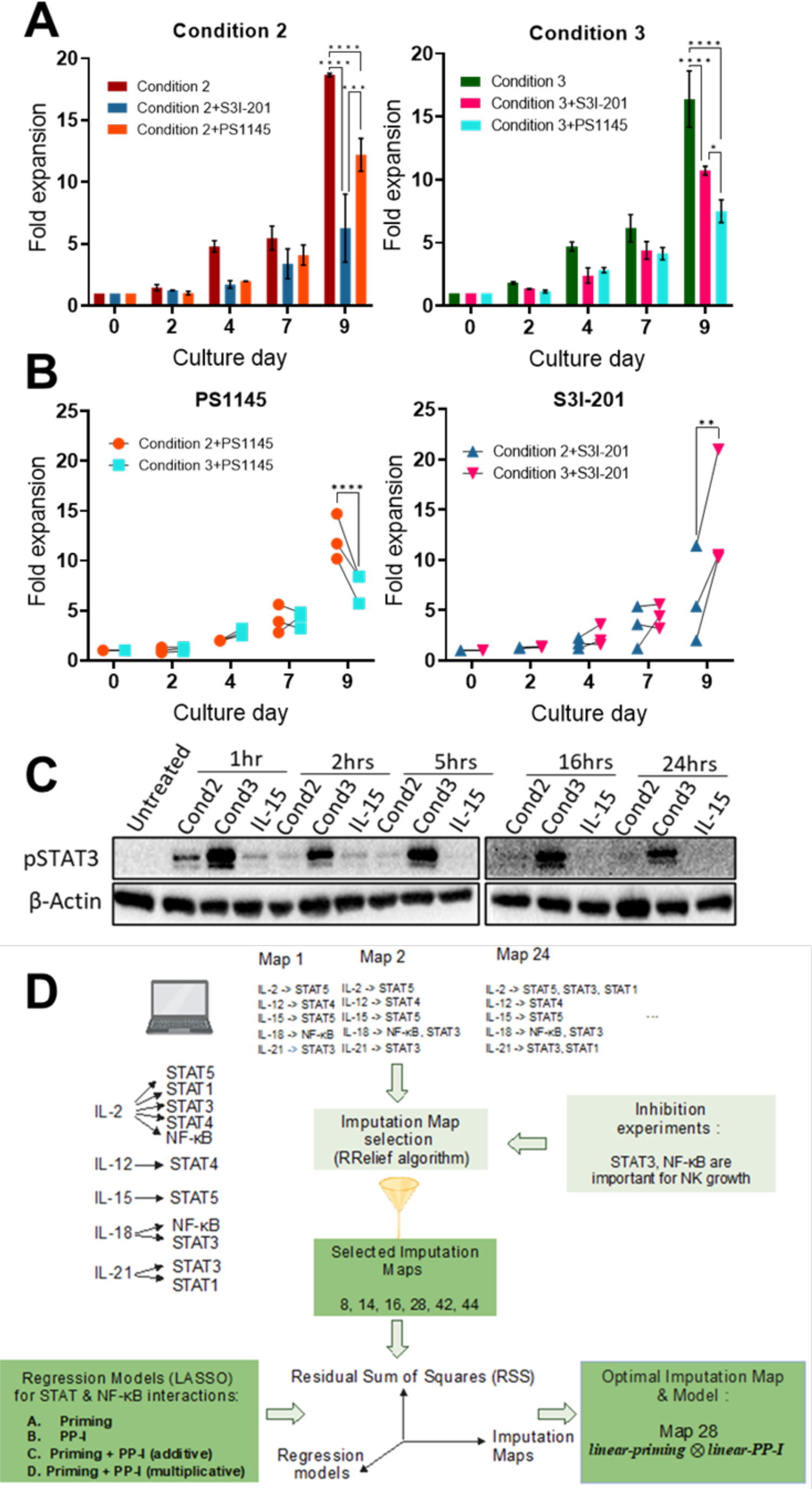
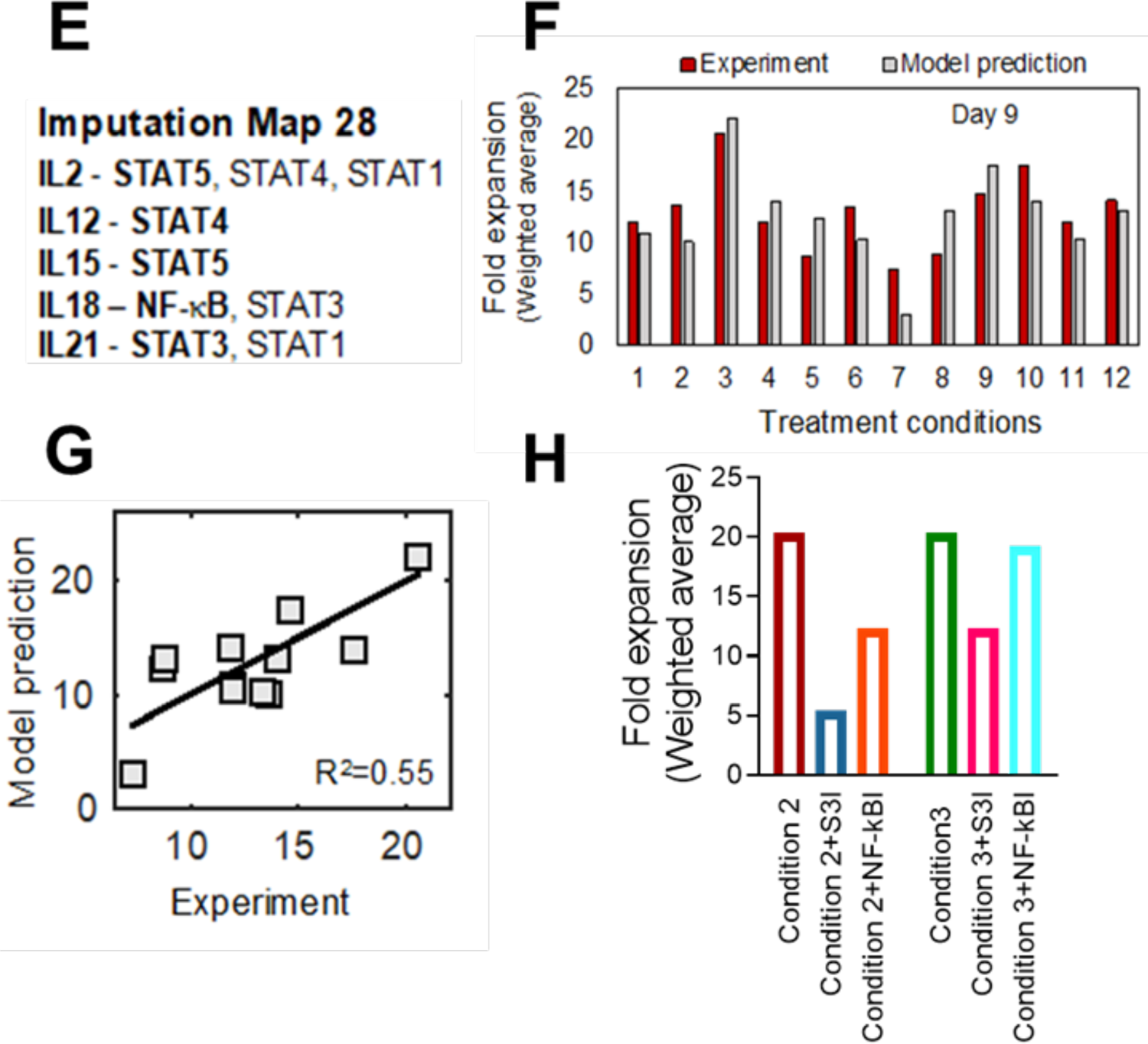
Developing and testing a model for NK cells proliferation: IL-21 sustains early proliferation in-vitro. **(A-B)** Isolated NK cells from 3 donors were cultured with the cytokine combination condition 2: IL-12+15+18 priming for 16 hours, then IL-2 only, and condition 3: IL-12+15+18+21 priming for 16 hours, then IL-2 only. During the priming stage, NK cells were treated either with 75 μM S3I-201 (STAT3 inhibitor) or 10 μM PS1145 (NF-κB inhibitor), cells were then washed and cultured only with IL-2. Data shown as mean ± SEM. The fold expansion for the two conditions is shown in (**A)**, and the comparison for the two conditions are shown in (**B)**. Data analysed by two-way ANOVA, and Tuckey’s multiple comparison test (*P≤0.05, **P≤0.01, ***P≤0.001, ****P≤0.0001 **(C)** Isolated NK cells were cultured with Condition 2, Condition 3 or IL-15 only (1ng/ml), at the indicated timepoints and then pSTAT3 and β-actin protein levels detected by immunoblotting. **(D)** Construction of the in silico predictive model for NK fold expansion for various cytokine cocktails. Multiple (n=64) imputation maps are created based on primary and secondary stats activated by IL-2, 12, 15, 18, 21 (**Table S2**). Applying a feature selection method (RRelief (27)) on the weighted average of NK fold expansion (**Table S1B**) on day 9, we chose only six imputation maps (map 8, 14, 16, 28, 42, 44) based upon experimental observations seen in inhibition experiments in Fig. 3A and **B**. Based on these six imputation maps (**Table S2**), multiple linear regression models are constructed considering various interactions of activated STATs and NF-κB between the priming and post-priming I periods across various cytokine cocktail conditions that regulate the NK cell fold expansion. To check the predictive ability of the models in estimating weighted average of fold expansion at day 9 in response to different cytokine cocktails, Leave-One-Out cross-validation (LOOCV) (26) is performed by varying imputation maps and linear regression models (see Materials and Methods). Optimal imputation map and regression model are chosen based on the minimum Residual Sum of Squares (RSS) across 12 LOOCV test sets. **(E)** STAT and NF-κB activation are shown for the optimal imputation map (Map 28) at day 9 gives the best predictive model. **(F)** Using the optimal imputation map and regression model, comparison of the fold expansion between experiment (maroon) and in silico predictive model (grey) are shown at day 9 for 12 cytokine cocktail conditions. **(G)** Overall prediction ability of weighted average of NK cell fold expansion (Fig. 3F) 12 cytokine cocktail conditions using the optimal map and model is shown at day 9. Model prediction vs. experiment in fold expansion gives R-square = 0.55 (see Materials and Methods for calculation of R-square). **(H)** Using optimal map and model, weighted average of fold expansion at day 9 are predicted for condition 2 and condition 3 with and without the presence of STAT3 and NF-κB inhibitors.

### Interactions between STATs and NF-κB factors induced by priming and post-priming cytokine stimulation regulate NK cell proliferation in vitro

Cytokine stimulation induces NK cells modulation by signaling via transcription factors such as STATs and NF-κB (**Table 1**). Experimental data has identified how specific cytokines pair with these receptor factors. However, many interactions are possible as STATs and NF-κB can be activated by multiple cytokines. Additionally, there is crosstalk between the STAT and NF-κB pathways which induce NK cell proliferation, with the nature of this crosstalk dependent on the cytokines used to stimulate NK cells. We created various possible imputation maps for our *in-silico* studies that explored potential hypotheses for primary and secondary STAT/NF-κB induced by specific interleukins (**Table S2)**. For example, IL-2 primarily activates STAT5, but can also activate STAT1, STAT3, STAT4, NF-κB as secondary STATs, whereas IL-18 primarily activates NF-κB but can also activate STAT3 and STAT1 as secondary STATs (**Table 1**). We considered the presence of activated forms of STATs and NF-κB which affect NK cell proliferation with binary variables (0 ≡ absent, 1≡ present). Subsequently, a feature selection procedure using an algorithm RRelief (27) was performed to determine a subset of maps (imputation maps 8, 14, 16, 28, 42, 44, **Fig. 3D** and **Table S2**) that are consistent with the experimental observation that STAT3 and NF-κB inhibition downregulates NK cell proliferation when cultured with both condition 2 and 3 (**Fig. 3A**). Next, we set up regression models to determine the effect of the crosstalk (synergistic or antagonistic) between the STATs and NF-κB activated by the cytokine stimulations during the priming and the post-priming I (PP-I) stages in NK cell proliferation (see Materials and Methods). We reasoned that activated STATs and NF-κB modify the rate at which single NK cells proliferate. Thus, we set up regression models where the weighted average of fold expansion of the NK cell population at a specific timepoint (e.g., day 9) with respect to its value in the unstimulated condition at day 0 is determined by the presence and absence of the STAT and NF-κB activation in the priming and the post priming stages (further details of weighted average in Materials and Methods, **Table S1B**). We trained or estimated the regression parameters for a set of cytokine conditions and then predicted the weighted average of fold expansion of NK cell population for a cytokine condition not included in the training (**Fig. 3D**). We considered four possible scenarios of STAT/NF-κB interactions that regulate the fold expansion of NK cell populations (see Materials and Methods, **Fig. 3D**): **(I)** STATs and NF-κB induced in the *priming phase* but not in the post-priming (PP-I) phase regulate NK cell proliferation. **(II)** STATs and NF-κB induced in the *PP-I phase* but not in the priming phase regulate NK cell proliferation. **(III)** STATs and NF-κB induced in the *priming- and post-priming (PP-I)* regulate NK cell proliferation *additively*. **(IV)** STATs and NF-κB induced in the *priming- and post-priming (PP-I) synergize or antagonize* and regulate NK cell proliferation (see Materials and Methods, **Fig. S1, Table S3**). We fitted the regression models with LASSO regularization (L1) (26) to the NK cell fold expansion data for each of the above scenarios. The NK cell fold expansion data show large variation across the donors containing outliers (“super donors”), that show very high NK cell expansion for a particular cytokine cocktail compared to other donors (see Donor 7 for condition 3 in **Table S1A**). Thus, we computed a weighted average of the fold expansion based on the variation in response of donors across different treatment conditions in the NK cell populations by using deviances (details in Materials and Methods section and **Table S1B**). The regression models for various scenarios and feasible maps are compared based on their prediction capabilities in terms of cross-validation error (26), where, to predict fold expansion for each observed cytokine cocktail condition or test data, we trained the regression model with the other observed 11 cytokine cocktail conditions in **Fig. 1C** by performing Leave-One-Out Cross-Validation (LOOCV). Finally, to determine the optimal imputation map and regression model, we evaluated their performance using a standard measure of overall prediction error: the Residual Sum of Squares (RSS) (26). This evaluation was conducted across 12 LOOCV test sets, with each set representing a different cytokine cocktail condition. For each regression model, R-square (see Materials and Methods) is calculated to evaluate overall predictions for those 12 observed conditions (**Table S3, Fig. S1**).

**Table 1.** List of activated STATs and NF-κB by ILs. The bold face and normal fonts represent the primary and the secondary STAT/NF-κB transcription factors, respectively.

| Cytokines | Induced STATs reported in literature | References | Induced STATs used in our model | Comments |
| --- | --- | --- | --- | --- |
| IL-2 | **STAT5**, STAT3, STAT4, STAT1, NF-κB | (6,40–42) | **STAT5**, STAT3, STAT1, STAT4 NF-κB | |
| IL-12 | **STAT4**, STAT3, STAT1, STAT5 | (6,40) | **STAT4** | For simplicity, we considered the primary STAT contribution alone |
| IL-15 | **STAT5**, STAT3, STAT1 | (6,40,43) | **STAT5** | For simplicity, we considered the primary STAT contribution alone |
| IL-18 | **NF-κB**, STAT3 | (44,45) | **NF-κB**, STAT3 |  |
| IL-21 | **STAT3**, STAT1, STAT5 | (6,40) | **STAT3**, STAT1 | Considered the primary and one secondary STAT contributions |

Our best predictive model shows imputation Map 28 (IL-2 → STAT5, STAT4, STAT1, IL-12→ STAT4, IL-15→ STAT5, IL-18→ NF-κB, STAT3, IL-21→ STAT3, STAT1) (**Fig. 3E**) among those selected feasible imputation maps with a strong pairwise synergy between the STAT3 in the priming and STAT1 in post-priming (PP-I) stimulation phases (*linear-Priming*⊗*linear-PP-I* in Materials and Methods and **Table S4**) is best suited (with the minimum RSS and the highest R-square, Materials and Methods, **Table S3** and **Fig. S1**) to describe fold expansion in NK cell populations (26). The optimum imputation map (**Fig. 3E**) and the synergy between STATs and NF-κB between priming and post-stimulation (PP-I) predicts that the cytokine condition 3 generates the maximum fold expansion compared to other conditions for typical donors which is validated in the experimental data (**Fig. 1C** and **Fig. 3F, Table S1B**). Next, we examined the predictive performance of the optimal model in determining the condition associated with the lowest fold expansion in the weighted average among the 12 conditions. Our model successfully identified condition 7 as the one yielding the minimum fold expansion, which is also validated in the experimental data (**Fig. 3F** and **Table S1B**).

Moreover, using the optimal map 28 and regression model (*linear-Priming*⊗*linear-PP-I)* we predicted weighted average of NK cell fold expansion (**Fig. 3H**) at day 9 for conditions 2 and 3 with and without the presence of STAT3 and NF-κB inhibitors. Both the STAT3-inhibitor (S3I) and the NF-κB-inhibitor (NF-κBI) reduce fold expansion in condition 2 in agreement with the *in vitro* data (**Fig. 3B**). Furthermore, the STAT3-inhibitor leads to a greater decrease in fold expansion than the NF-κB-inhibitor for condition 2, reproducing the fold expansion behavior observed in experiments (**Fig. 3B**). For condition 3 the STAT3-inhibitor in the model shows a reduced fold expansion, in agreement with experimental in vitro observations (**Fig. 3A**). However, the reduction in the fold change in condition 3 is not strong enough for NF-κB-inhibitor and has lesser effect than STAT3-inhibitor in reducing fold expansion indicating a limitation in the training of our model.

Despite this limitation, our model provides valuable quantitative insights into the contributions of STAT3, NF-κB, and other induced STATs during the priming and post-priming phases. It also sheds a light on the intricate interplay and potential synergistic or antagonistic interactions among these factors in driving the proliferation of NK cells under the influence of different cytokine cocktails (**Table S4**). This suggests that condition 3 is an optimal culture condition, where the presence of IL-21 yields the highest NK cell fold expansion in weighted average at day 9. Conversely, condition 7, lacking IL-21 in the priming phase and IL-15 in the post-priming phase, exhibits the lowest NK cell fold expansion in weighted average (**Fig. 3F** and **Table S1B**).

### IL-21 does not impact cytotoxicity

Successful immunotherapy depends on NK cell functionality as well as NK cell numbers, therefore, to determine how the different cytokine combinations affected NK cell effector functions we studied degranulation (CD107a) and interferon gamma (IFN-γ) production using NK cells cultured under different protocols with and without IL-21 (**Fig. 4A** and **Fig. 4B**). These were then tested in co-culture assays with three different hepatocellular carcinoma (HCC) cell lines at day 10. The introduction of IL-21 in the priming phase did not affect cytotoxicity or IFN-γ secretion (condition 2 versus condition 3). Overall effector function was most notable for condition 12 which gave the strongest results for both degranulation and IFN-γ secretion against the cell lines tested. This regimen contains IL-21 in the priming phase and IL-18 in the post-priming phase, and strong cytotoxicity was seen against two out of the three cell lines tested, but strong IFN-γ secretion against all the cell lines tested (**Fig. 4A** and **Fig. 4B**). Thus, it appears that the dominant effect on cytotoxicity was in having IL-18 in the post priming phase.

**Figure 4.**
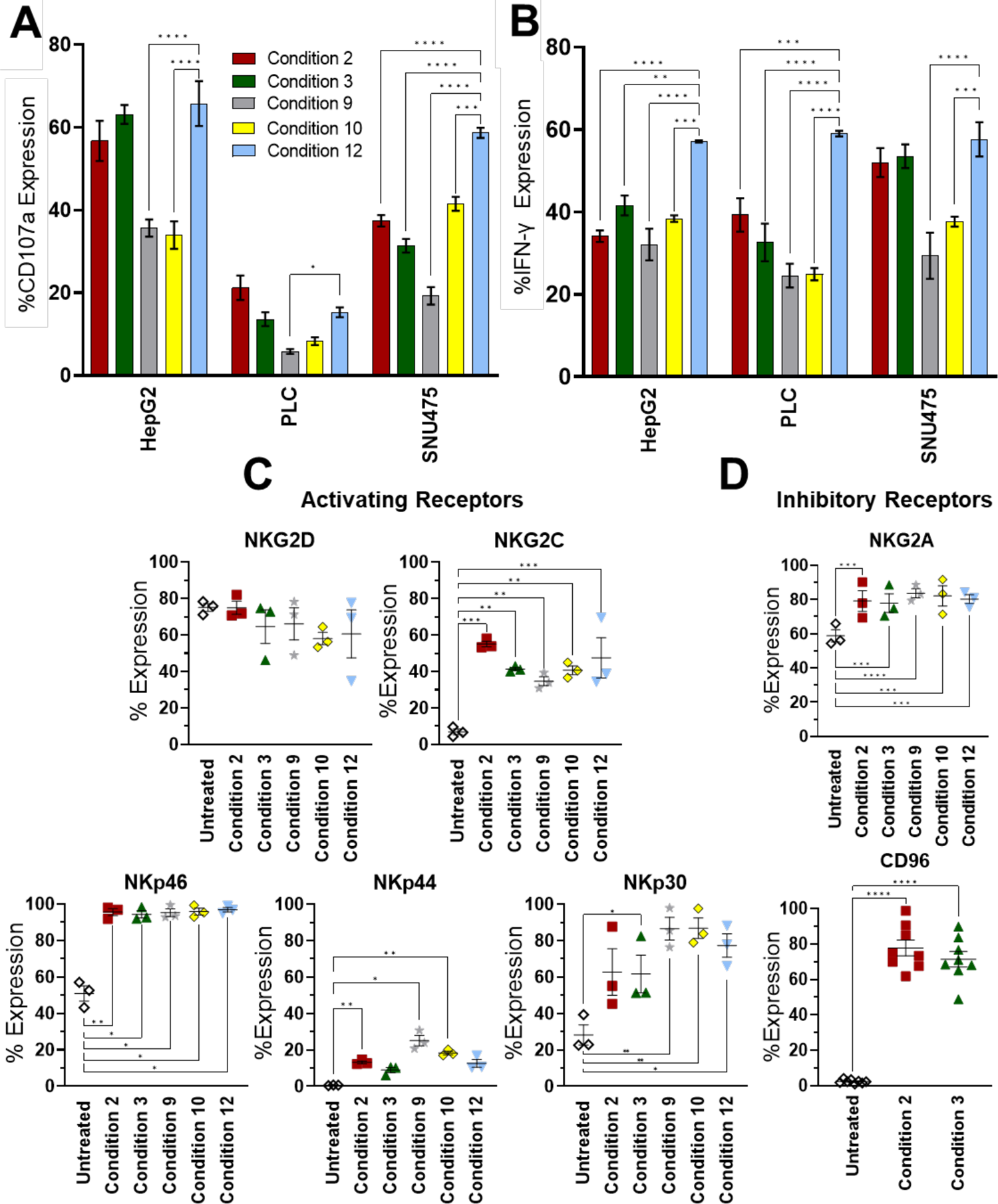
Effects of different regiments on NK cell receptor expression and cytotoxicity against hepatocellular carcinoma cell lines as a model for immunotherapy. NK cells from 3 donors were cultured using the indicated cytokine conditions and then assayed on day 10 for cytotoxicity against the indicated hepatocellular carcinoma cell lines using a CD107a degranulation assay (**A**) or for expression of IFN-γ (**B**). Assays were performed at an effector, target ratio of 1:1. Data analysed by two-way ANOVA, and Dunnett’s multiple comparison test (*P≤0.05, **P≤0.01, ***P≤0.001, ****P≤0.0001). **(C)** Expression of activating receptors by NK cells stimulated by cytokines for 10 days. **(D)** Expression of inhibitory receptors by NK cells stimulated by cytokines for 10 days. Data shown as mean± SEM (n=3), analysed by RM one-way ANOVA, and Dunnett’s multiple comparison test (*P≤0.05, **P≤0.01, ***P≤0.001, ****P≤0.0001).

To correlate the effector functions with cytokine conditions we studied receptor expression. Overall, there were no simple correlations of receptor expressions with cytokine condition. In particular, expression of the inhibitory receptor NKG2A was high for all regimens tested and of the activating receptors NKG2D, NKG2C and NKp46 were similar amongst the different regimens (**Fig. 4C**). There was a trend towards lower NKp44 and NKp30 in the conditions associated with IL-12/15/18 priming.

To compare the cytotoxic responses of NK cells against HCC cell lines, we measured CD107a expressions against HepG2, PLC, SNU475 and Huh7 for conditions 1-6. We observed the highest cytotoxic responses against HepG2 and Huh7 cell, with a moderate response against SNU475 and minimal response against PLC (**Fig. 5A**). To investigate how receptor expression determined cytotoxicity of these lines, we performed correlation analysis to evaluate Pearson correlation coefficients between expressions of CD107a and activating/inhibitory receptors (**Fig. S2, S3, S4** and **Table S5**). However, it did not generate any positive correlations of cytotoxicity with expression of any specific activating receptor for these cell lines except Huh7. We therefore assessed cytotoxicity of NK cells (8 donors, D3-D10) against Huh7 cells using a different combination of cytokines conditions 1-6, which allowed assessment of the use of IL-21 in the priming (condition 3) and post-priming periods (condition 6) (**Fig. 5A**). We found that CD107a expression on CD56+CD16+ cytotoxic NK cells had a positive correlation with levels of the activating receptor NKp44 (ρ=0.71, P=9.3×10^-5^) for the Huh7 cell line but there were no other positive correlations noted with the other receptors (**Fig. 5B**). Conversely, there was a negative correlation with the inhibitory receptor NKG2A (ρ= −0.63, P= 9.1×10^-4^) (**Fig. 5B**). This is extended to the other cell lines (HepG2, PLC and SNU475) (**Fig. 5C**). Thus, using multiple different NK cell expansion protocols, the dominant determinant in killing HCC cell lines appears to be NKG2A expression. To evaluate the impact of presence of IL-21 and IL-18 in cytokine conditions on donors, we grouped the conditions (**Fig. 5B, 5C**), which shows that donors treated with IL-21 or others (IL-2, IL-15) shows higher cytotoxicity against Huh7 cell line (**Fig. 5B**). We further observed that within these NK cell expansion protocols, the activating receptor NKG2C, expressed in memory NK cells, correlated positively with the inhibitory receptor NKG2A (ρ=0.854, P=2.8×10^-10^) and also there were positive correlations between the following pairs of activating receptors: NKp44 and NKp30 (Pearson correlation, ρ=0.891, P=3.5×10^-12^), and NKp46 and NKG2C (ρ=0.563, P=6.5×10^-4^), and negative correlations are present between NKG2D and NKp30 (ρ= −0.619, P=1.2×10^-4^) and, NKG2D and NKp44 (ρ = −0.696, P=6.9×10^-6^) (**Fig. 5D**).

**Figure 5.**
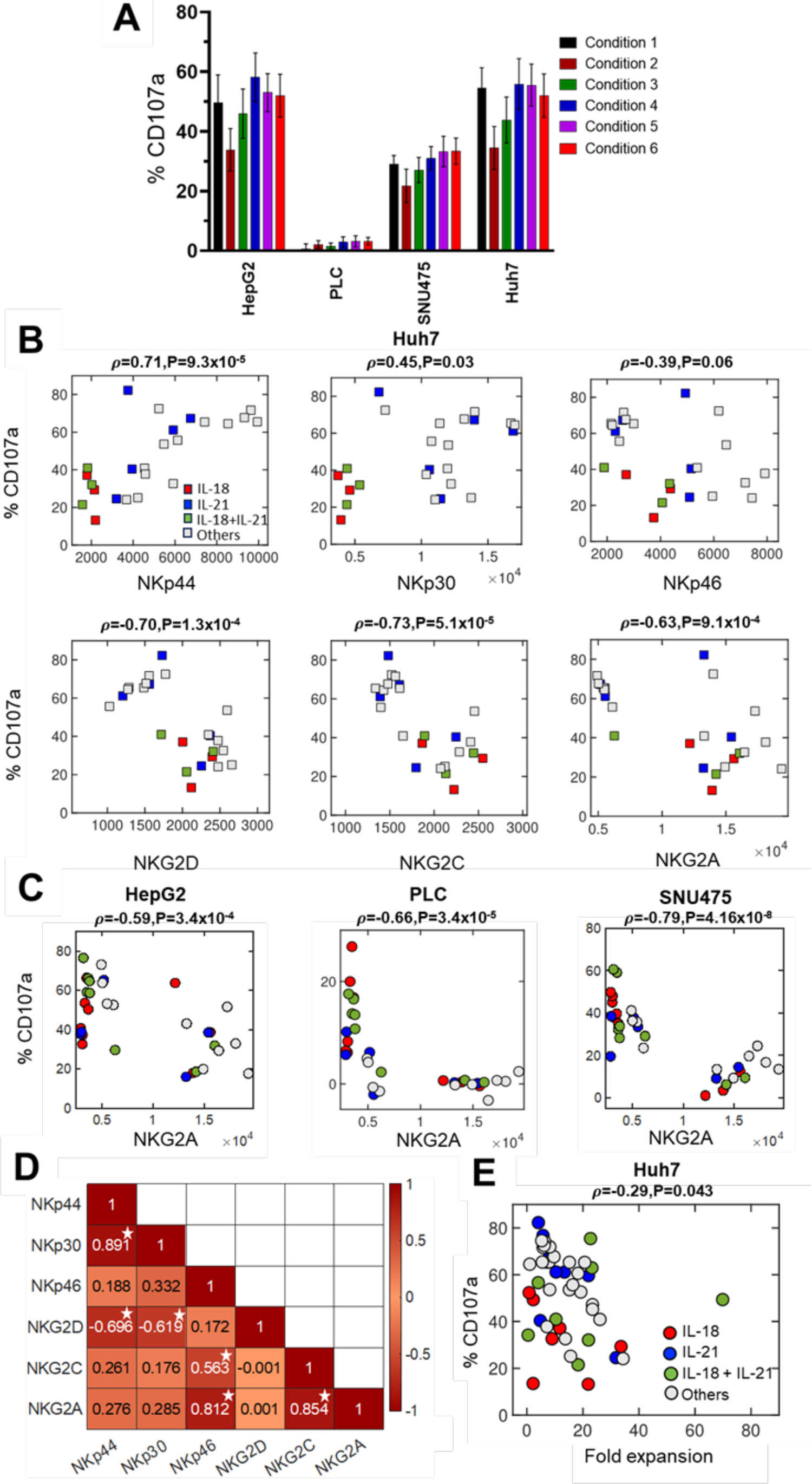
Correlations of cytotoxicity with NK cell proliferation and receptor expressions against different hepatocellular carcinoma cell lines. (**A**) % CD107a expression levels against four hepatocellular carcinoma cell lines HepG2, PLC, SNU475 and Huh7. NK cells from 8 donors (donor 3 to 10) donors were incubated with cytokines according to conditions 1-6 and then assayed for proliferation, CD107a and receptor expression on Day 10. (**B**) Correlation of cytotoxicity (%CD107a expression) of the Huh7 cell line with NK cell receptor mean expressions for the high CD16 expressing donors (8 donors, D5-D6, D8-D10, D15-D17) cultured under 10 different cytokine conditions (conditions 1-6, 9-12). Each circle represents a donor, labelled based on the treatment conditions in the presence of either IL-18 (red) or, IL-21 (blue) or, both IL-18+21 (green) or none (gray). Pearson correlations (ρ) and *P* values are shown between %CD107a and each of mean receptor expressions. (**C**) Correlation of NKG2A expression on NK cells with cytotoxicity for HepG2, PLC and SNU475 cell lines showing Pearson correlations (ρ) and *P* values from high CD16 expressing donors. (**D**) Correlation matrix representing the correlation between the mean receptor expressions on the NK cells from high CD16 expressing donors. Pearson correlation coefficient values are shown in each box. A darker color indicates higher correlation (positive or negative) between the two receptors. Boxes with white star indicates Pearson correlation values with P < 0.05. **(E)** Against Huh7 cell line, %CD107a expression from (**A)** was correlated with average of NK cell fold expansion at day 9 (Fig. 1C) for 8 donors (D3–D10), each treated with conditions 1-6.

Finally, we determined how proliferation impacted cytotoxicity. For the Huh7 cell line, overall using the 6 different regimes (conditions 1-6) tested in 8 donors (D3-D10) we observed that there was generally a negative impact of proliferation on cytotoxicity (ρ=-0.29, P=0.043), with one single donor responding having both strong proliferation and cytotoxicity (**Fig. 5E**, Donor 7 for condition 3 in Table S1A). However, there was no specific segregation of regimens using the different cytokine combinations (other cell lines in **Fig. S2, S3, S4**).

## Discussion

Characterization and profiling of NK cells is essential to better understand their biology, so they can be modulated for future NK cell-based immunotherapy. There is a complex interplay between cytokines that dictate the functions of NK cells, and which needs to be understood in order to identify better combinations of cytokines for clinical use. We addressed this challenge by developing an *in-silico* framework to predict fold expansion in populations of NK cells treated with cytokine cocktails. Our results indicated that NK cell proliferation can be modulated during both the short 16 hours priming and a 3-day post-priming period. Our modeling suggested the importance of the synergy between STAT3 induced by IL-21 in the initial priming period and that this interacts with the other STATs such as STAT1 in post-priming period in increasing the NK cell proliferation (**Table S4**). Previous studies show roles of STAT3 in conjunction with STAT5 in suppressing NK cell cytotoxicity (6), however, our experimental and modeling studies demonstrate a synergistic role of STAT3 in increasing NK cell proliferation.

We focused primarily on IL-21, as this has been used predominantly in a membrane bound form to induce proliferation of NK cells, but we sought to identify opportunities for using this cytokine in its physiological soluble form in order to investigate its potential without the use of feeder cells. IL-21 induces STAT3 activation leading to downstream activation of c-Myc which regulates cellular processes such as increasing NK cell proliferation, induction of glycolysis, mitochondrial biogenesis, and the cell cycle (28). NK cells with a high STAT3 expression following IL-21 stimulation have rapid proliferation, long telomeres, and resistance to senescence (28). This may be important in making NK cells more resistant to immunosuppressive tumor microenvironments and hence enhance their activity against “cold” tumors, such as HCC (29–32). Solid tumors are less susceptible to NK cell therapies as compared to hematological tumors and therefore making NK cells less susceptible to these immunosuppressive effects will be key to generating successful NK cell-based treatments for this class of cancers. Although IL-21 signals via a combination of STATs, NF-κB inhibition caused a lower proliferation in the presence of IL-21, indicative of the complex interplay between these two pathways on NK cell proliferation, which has previously been observed in cancer (33). The importance of IL-21 and NF-κB in mediating NK cell expansion is highlighted by the significant reduction of NKG2A levels following NF-κB inhibition, with expression decreasing from approximately 70% to ∼35% in the presence of IL-21 (condition 3, Supplementary **Fig. S6B**). This is in line with *Kaulfuss et al* (34), who showed that NK cells expansion was greater in NKG2A^+^ NK cells and that this was not dependent on STAT3 signaling pathway. This is consistent with our data showing the tradeoff between proliferation and killing, and the negative effects of NKG2A expression on all HCC targets tested.

A major challenge in predicting changes in population sizes of NK cells treated with cytokine cocktails is the differences in the induction of phosphorylated forms of STAT and NF-κB factors which can potentially involve a large number (∼100) of combinations of STAT and NF-κB factors that can synergize/antagonize to affecting NK cell proliferation. In the cytosol, the STATs form homo- and hetero-dimers, and in certain cases tetramers, where the formation of specific multimers can depend on the concentrations of phosphorylated forms of STATs generated during signaling (35–37). The STAT dimers can further synergize/antagonize in the nucleus where these might competitively/co-operatively engage with DNA regulatory elements and affect co-operative gene regulatory processes (12,38,39). We confronted this challenge in large dimensions by implementing a feature selection algorithm RRelief for fold expansion of NK cell populations in treatment conditions containing different combinations of interleukins and repeating some of these experiments with STAT3 and NF-κB inhibitors. The combined approach allowed us to select a small number of potential possibilities for STAT and NF-κB induction in the cytokine cocktails that are consistent with the inhibition experiments (**Fig. 3A**). The next challenge of relating the phosphorylation of STAT and NF-κB factors, their synergy/antagonism, and their influence on proliferation was addressed by using a linear regression model with LASSO regularization. Cross-validation of the possible models against the NK cell fold expansion data revealed that the synergy/antagonism between the factors induced in the priming and post-priming play an important role in affecting NK cell proliferation. The success of our framework in predicting the cytokine cocktail with the largest fold expansion demonstrates the utility of such combined approach in modeling NK cell response against cytokine cocktails. This methodology, initially studied at day 9, holds promise for further investigation at other days of proliferation, and perhaps for modeling maturation of NK cells in cytokine cocktails. Our work integrates NK cell proliferation, function, and phenotype to develop a model for NK cell generation that supports the incorporation of IL-21 into short term NK cell proliferation regimens. The next step would be to incorporate these into pre-clinical models for cancer immunotherapy.

### Limitations of the study

The main limitation of the biological aspects of this study are the use of multiple different donors, and not all were tested with the same regimes. This was due to sample volume limitations; purified NK cells were used for these assays. However, this limitation may also be considered a strength of the work, as it is important to sample different donors to generate a widely model that is widely applicable to the population. This is because it is known that there is substantial donor-to-donor variability, which affects NK cell proliferation. It should also be noted that there was a substantial donor-to-donor variability which is a known feature of NK cell proliferation. Our regression models did not consider dependencies of NK cell proliferation on cytokine concentrations and subcellular concentrations of the induced STAT and NF-κB proteins which could further regulate synergy/antagonism between these transcription factors. In this study, our model framework was employed to predict NK proliferation at day 9, which can be extended to study other days. However, the optimal map and regression model could be different at other days than day 9. For our optimal predictive model selection, the imputation map and regression model representing STAT/NF-κB interactions chosen in our study, might not uncover the true mechanism, yet serve as a crucial mathematical framework for generating inferences in the context of cytokine conditions.

## Material and Methods

### Peripheral Blood Mononuclear Cell (PBMC) isolation and NK cell isolation

Peripheral blood mononuclear cells (PBMC) were isolated from buffy cone of healthy donors by density gradient centrifugation, using Ficoll-Hypaque (Fisher Scientific UK) following the manufacturer instructions. CD56+CD3-NK cells were purified from PBMC, by positive selection by using the Miltenyi NK isolation kit with LS columns (Miltenyi Biotec, Surrey, UK), as stated by the manufacturers’ protocol.

### Cell lines

Liver cancer cell lines, HepG2, PLC cultured in complete DMEM, 10% fetal bovine albumin (Sigma-Aldrich, UK), and 5% PenStrep (Life technologies, UK); the liver cancer cell line SNU475 was cultured in complete RPMI, supplied with 10% fetal bovine albumin and 5% PenStrep. Cells were kept in a humidified incubator at 37°C and 5% CO_2_.

### NK cells stimulation

NK cells were maintained in culture in NK MACS medium (Miltenyi, UK), supplemented with 1% NK supplements (Miltenyi, UK), 1% PenStrep (Life technologies, UK); 5% human AB serum (Sigma-Aldrich, UK). NK cells were expanded with cytokines IL-2 (500 U/ml), IL-12 (10 ng/ml), IL-15 (20 ng/l), IL-18 (50ng/ml), IL-21 (25 ng/ml) (R&D System, UK), for 10 days.

### NK cells staining

NK cells were analyzed for receptor expression using the following antibodies, CD56 (BV510, Biolegend), CD3 (APC/Cy7, Biolegend), NKp30 (PerCp/Cy5.5, Biolegend), NKG2A (FITC, Miltenyi), NKG2C (PE, Miltenyi), NKG2D (FITC, Biolegend), CD94 (BV421, Biolegend), NKp44 ((PerCp/Cy5.5, Biolegend), NKp46 (APC, Biolegend), CD16 (BV421, Biolegend), CD57 (APC, Biolegend), CD117 (PE/Cy7, Biolegend), CD16 (BV421, Biolegend). For CFSE staining unstimulated NK Cells were stained with CellTrace CFSE (Thermo Fisher Scientific, UK), according to manufacturer’s instructions and assessed for staining by flow cytometry. Data were analyzed using FlowJo v.10.9.0.

### Degranulation assay and assessment of IFN-γ expression

NK cells were cultured with cell line in complete RPMI medium, with anti-human CD107a-eFluor660 (clone H4A3, Invitrogen, UK) at an E:T ratio of 1:1 for a total of 4 hours at 37°C. After 1 hour of incubation, GolgiStop Protein Transport Inhibitor (BD Bioscience, UK) was added, and cells were incubated for a further three hours. Cells were then stained with antibodies CD56 (PE/Cy7, Biolegend), CD3 (BV510, Biolegend) and IFN-γ (PE, Biolegend) and analyzed by flow cytometry. Gating strategy shown in Fig. **S5.**

### Model development

#### Terminology for priming and post-priming (I & II)

The experimental procedure consists of three distinct stages: priming, post-priming I & post-priming II (Fig. 1B). During the priming stage, NK cells are exposed to specific cytokine cocktails for a period of 16 hours. This initial treatment is designed to activate and prepare the NK cells for subsequent analysis. After the priming stage, the culture media is replaced, and the cells are subjected to a new set of cytokine cocktails. This second phase, lasting from 16 hours to 3 days, is referred to as the post-priming I (PP-I) period (Fig. 1B). Media is again replaced at day 3 and new set of cytokines are added for day 3 to day 16, which is referred as the post-priming II (PP-II) period.

#### Calculating Weighted average (Y) for NK cell fold expansion

The NK proliferation data shows high variability in fold expansion across donors. Thus, to choose a better representative quantity, instead of calculating the standard average of fold expansions across donors (Fig. 1C), we calculated the weighted average for the fold expansion (Fig. 3F and **Table S1B**). Here, we considered unequal weights for individual donors based on his/her responses in fold expansion across various cytokine cocktail conditions. For example, if NK cells from a donor are treated with *k* cytokine cocktail conditions (e.g., conditions 1-6, i.e., *k*=6), we calculated the variances (σ^2^ = *var*({*f*_i_^1^, *f*_i_^2^, …, *f*_i_^k^}) of NK cell fold expansion (*f*_i_*^α^*) across *α* =1,…,*k* cytokine conditions for that *i*^th^ donor. The data shows that different donors have small or, moderate, or high variances in fold expansion across conditions. We assigned significant weights for donors with moderate variances in fold expansion across cytokine cocktail conditions and less for donors with very high or low variances. In such a way, we assigned small weights to the outliers, either non-responders (low variance in NK cell fold expansion across *k* cytokine conditions) or super-responders (very high variance in NK cell fold expansion across *k* cytokine conditions), and considered the responses significantly from the typical responders around median of variances (*median*({σ_1_^2^, σ_2_^2^, …, σ_N_^2^})) in fold expansion from *N* donors. This is systematically done by choosing the weights for the individual donor (*ω*_i_) as inversely proportional to their deviance 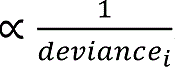. By normalizing the weights, we get the weight for *i*^th^ donor as 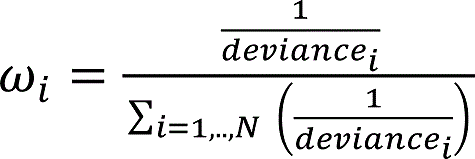, such that Σ_i=1,..,N_ω_i_=1. Where deviance of i^th^ donor, *d*_i_ = (σ_i_^2^ − *medme*)^2^, where *medme* = [0.1 × *mean*({σ_1_^2^, σ_2_^2^, …, σ_N_^2^}) + 0.9 × *median*({σ_1_^2^, σ_2_^2^, …, σ_N_^2^})]. The weighted average of fold expansion for *α*^th^ cytokine condition is calculated as Ŷ*^α^* = ∑^%^ *α*^th^ cytokine condition, Where *f*^α^_i_ represents the fold expansion for *i*^th^ donor treated with α^th^ cytokine condition.

In our data (**Table S1A**), since all 17 donors are not treated with same cytokine conditions, we divided them into three groups: (I) Group A – *N_A_* (=10) donors treated with conditions 1-6 (Donor 1-donor 10), (II) Group B – *N_B_* (=4) donors treated with conditions 7-8 (Donor 11-Donor 14) and (III) Group C – *N_C_* (=3) donors treated with conditions 9-12 (Donor 15-Donor 17) and calculated weighted average of fold expansion and for *α*^th^ cytokine condition for group A, B and C, respectively. Then we combined them to calculate weighted average of fold expansion *Y* = {*Y*_A_, *Y*_B_, *Y*_C_} across 12 conditions.

#### Generation of Imputation Maps: IL to STAT/NF-κB conversion

Cytokines in a cytokine cocktail can show synergistic or antagonistic effects in NK cell responses (proliferation or degranulation). Each cytokine activates multiple STATs, which include a primary STAT along with secondary STATs (**Table 1**). Since the type of activated STATs by a particular cytokine is limited (seven), the STATs and NF-κB induced by different cytokines can overlap. Consequently, different cytokine cocktails can activate the same sets of STATs, albeit in different concentrations. IL-2 and IL-15 activate primarily STAT5, whereas IL-12, IL-18, and IL-21 primarily activate STAT4, NF-κB and STAT3, respectively. Additionally, IL-2 activates STAT1, STAT3, STAT4, NF-κB as secondary STATs. Whereas IL-18 and IL-21 activate STAT3 and STAT1 respectively as secondary STATs.

In our modeling framework, we considered the presence or the absence of STATs in a binary representation (0 or, 1) due to the lack of knowledge about the true concentrations of activated STATs for a specific cytokine cocktail. In addition, it is unclear what set of secondary STATs is activated in by a particular cytokine cocktail. Therefore, we assumed in our model that each interleukin in a cytokine cocktail activates either a primary STAT alone or a primary STAT along with a set of secondary STATs and NF-κB chosen from all possible STAT and NF-κB combinations (single or multiple) (**Table 1**). For example, IL-2 can induce STAT5 as a primary STAT alone or STAT3, STAT4, STAT1 and NF-κB as secondary STAT/NF-κB along with it in 16 (=1+^4^C_1_+^4^C_2_+^4^C_3_+^4^C_4_) ways. Similarly, IL-12, IL-15, IL-18, and IL-21 can induce secondary STATs and NF-κB in 1, 1, 2 and 2 ways, respectively. Considering all the possible combinations of activated STATs and NF-κB in a cytokine cocktail, there can be up to 16×1×1×2×2 = 64 possible sets (indexed by *i*) of primary and secondary STATs (Fig. 3D and **Table S2**). Each of these possibilities represents a specific imputation map (indexed by *i*) which generates activated STATs and NF-κB as the outputs for the cytokine combinations used in the treatment conditions. We create a binary matrix, **M***_i_*(*p,q*) of size 12×10, where the rows (*p*) and the columns (*q*) represent the cytokine conditions and the STATs and NF-κB, respectively, and the matrix elements describe the presence of absence of STATs and NF-κB for any cytokine condition given by the imputation map *i*. Columns 1-5 and 6-10 represent the presence or absence of STAT1, STAT3, STAT4, STAT5, NF-κB during priming and PP-I, respectively. Thus, applying these possible 64 imputation maps to cytokine treatment conditions we constructed 64 binary matrices **M***_i_*, where, *i*=1,…,64.

#### Reduction of Imputation Maps based on Inhibition experiments in Fig. 3(A)

We selected imputation maps that can describe the observations with STAT3 and NF-κB inhibitors in the following way. We imposed two constraints observed in the experiments with the inhibitors (See Fig. 3A). *Constraint 1:* The STAT3 inhibitor in condition 2 significantly decreases the NK cell proliferation fold expansion. Thus, we imposed a constraint that condition 2 must induce STAT3. *Constraint 2:* STAT3 and NF-κB inhibitors significantly decrease the NK cell proliferation fold expansion for conditions 2 and 3. Therefore, we impose the constraint that the presence of STAT3 and NF-κB increase NK cell proliferation. We evaluated the contribution of each STATs and NF-κB during priming and PP-I to NK cell fold expansion at day 9 in terms of RRelief weights (or scores) in the range of [-1,+1] using a feature selection method RRelief (27). Using the weighted average of NK cell fold expansion at day 9 (***Y***) (**Table S1B**), we first obtained the weight (or score) of each STATs/NF-κB during priming and PP-I for an imputation map *i* as a vector ***W_i_***. Imposing the two constraints above, we selected those maps for which ***W_i_*** contains positive weights corresponding to STAT3 and NF-κB both in priming and PP-I (Fig. 3D) to ensure STAT3 and NF-κB arise as essential feature variables in regulating NK cell proliferation in the feasible imputation maps. Using this filtering technique, we reduced the number of feasible imputation maps from 64 to 6 that are consistent with experimental observations (Fig. 3D).

#### Multiple linear-regression model to predict weighted average (Y) of NK fold expansion for a cytokine cocktail condition at certain day

To train and predict weighted average of fold expansions at day 9 of NK cell populations under different treatment conditions, represented by a vector ***Y*** (**Table S1B**), we constructed multiple linear regression models ***Y***=**X*β***+ɛ (26). Where, **X** represents a binary matrix constructed considering the possible interactions between the STATs and NF-κB in priming and PP-I. In our notation each element (indexed by α) of the ***Y*** vector represents a treatment condition. We considered additive or additive and pair-wise synergistic or antagonistic contributions of the induced STATs and NF-κB in the priming and PP-I in regulating NK cell fold expansion. The STATs and NF-κB are represented by binary vectors ***S_p_*** where a STAT or NF-κB (indexed by *p*) is present (=1) of absent (=0) in treatment condition given by the elements of the vector ***S*_p_**. The contribution of the STATs and NF-κB, represented by ***S_p_^(prime)^*** and ***S_p_^(PP-I)^***, in the priming and PP-I phase to the fold expansion at day 9 is quantified by C_α_^(prime)^=∑_p_ [*a*_p_[S_p_^(prime)^]_α_ + ∑_q,p≠q_ *b*_pq_[S_p_^(prime)^]_α_[S_q_^(prime)^]_α_] and C_α_^(PP-I)^=∑_p_ [*c*_p_[S_p_^(PP-I)^]_α_ + ∑_q,p≠q_ *d*_pq_[S_p_^(PP-I)^]_α_[S_q_^(PP-I)^]_α_], respectively. The coefficients b_pq_ and d_pq_ are set to zero if no synergy/antagonism is considered within the priming and post-priming phases. We considered four scenarios (8 regression models, see **Fig. S1** and **Table S4**) for combining STAT and NF-κB contributions in the priming and post-priming phases to describe the weighted average of NK cell fold expansion in NK cell proliferation.

A. ***Priming alone*:** In this model the fold expansion in NK cell proliferation is determined by the priming alone, and ***Y***=***C^(prime)^***. We consider two possible ways to include the STAT/NF-κB contributions. (i) *Linear contribution in priming*(*linear-priming*): Here the contributions of STAT/NF-κB arise as a sum of linear terms of {S_p_} such that *Y_α_*=∑_p_ [*a*_p_[S_p_^(prime)^]_α_, where the index α represents the cytokine cocktail condition. This leads to 5 predictor variables in design matrix **X** in presence of activated STAT1 (S1), STAT3 (S3), STAT4 (S4), STAT5 (S5) and NF-κB (Sb). (ii) *Linear + pairwise contribution* (*linear+pairwise-priming*): The contribution is constructed with the linear and pairwise STAT and NF-κB terms, present in the priming stage with the presence of S1, S3, S4, S5, Sb and their all possible cross terms*: Y_α_* =∑_p_ [*a*_p_[S_p_^(prime)^]_α_ + ∑_q,p≠q_ *b*_pq_[S_p_^(prime)^]_α_[S_q_^(prime)^]_α_]. This leads to 15 predictor variables in the design matrix **X**.
B. ***Post priming alone***: Here the fold expansion in NK cell is determined by the post-priming (PP-I) alone, and ***Y=C^(PP-^*^I)^**. We consider two models. (i) *Linear contribution (linear-PP-I):* Contains sum of linear STAT/NF-κB terms, i.e., *Y_α_=*∑_p_ *c*_p_[S_p_^(PP-I)^]_α_. This leads to 5 predictor variables in the design matrix **X**. (ii) *Linear + pairwise contribution (linear+pairwise-PP-I):* Contains linear and pairwise STAT/NF-κB terms. Thus, *Y_α_=*∑_p_ [*c*_p_[S_p_^(PP-I)^]_α_ +∑_q,p≠q_ *d*_pq_[S_p_^(PP-I)^]_α_[S_q_^(PP-I)^]_α_]. This leads to 15 predictor variables in the design matrix **X**.
C. ***Additive contribution***: In this case the fold expansion NK cell proliferation is given by ***Y=C^(prime)^ + C^(PP-I)^***. We consider two models. (i) *Linear contributions from the priming and the PP-I stage (linear-priming* ⊕ *linear-PP-I):* Here the regression is given by *Y_α_* =∑_p_ [*a*_p_[S_p_^(prime)^]_α_+ *c*_p_[S_p_^(PP-I)^]_α_]. This leads to 10 predictor variables in the design matrix **X**. (ii) *Linear + Pairwise contributions from the priming and the PP-I stage (linear+pairwise-priming* ⊕ *linear+pairwise-PP-I):* The regression model is given by, *Y_α_*=∑_p_ [*a*_p_[S_p_^(prime)^]_α_ + ∑_q,p≠q_ *b*_pq_[S_p_^(prime)^]_α_[S_q_^(prime)^]_α_] *+* ∑_p_ [*c*_p_[S_p_^(PP-I)^]_α_ + ∑_q,p≠q_ *d*_pq_[S_p_^(PP-I)^]_α_[S_q_^(PP-I)^]_α_]. This leads to 30 predictor variables in the design matrix **X**.
D. ***Multiplicative contribution***: In this case the fold expansion NK cell proliferation is given by, *Y_α_* =C^(prime)^_α_C^(PP-I)^_α_. We consider two models. (i) *Pairwise contributions from the priming and the PP-I stage (linear-priming* ⊗ *linear-PP-I):* Here the regression model is given by, *Y_α_=*(∑_p_ *a*_p_[S_p_^(prime)^]_α_)× (∑_p_ *c*_p_[S_p_^(PP-I)^]_α_) This leads to 25 predictor variables in the design matrix **X.** (ii) *Pairwise + higher order contributions from the priming and the PP-I period (linear+pairwise-priming* ⊗ *linear+pairwise-PP-I):* The regression here is given by, *Y_α_*= (∑_p_ [*a*_p_[S_p_^(prime)^]_α_ + ∑_q,p≠q_ *b*_pq_[S_p_^(prime)^]_α_[S_q_^(prime)^]_α_]) × (∑_p_[*c*_p_[S_p_^(PP-I)^]_α_ + ∑_q,p≠q_ *d*_pq_[S_p_^(PP-I)^]_α_[S_q_^(PP-I)^]_α_]). This leads to 225 predictor variables in the design matrix **X**.

We group unique STAT/NF-κB contributions, represented by unique terms in the right-hand sides of the above regressions, by the parameters β_k_ and explanatory (predictor) binary variables X_k_, denoting S_p_ or product of several S_p_, and estimate the parameters β_k_ using LASSO regression (**Table S4**). *β* > 0 represents the synergy between pair-wise STATs and NF-κB, and *β* < 0 illustrates antagonism.

#### Calculating the linear regression coefficients (β) to quantify synergy or antagonism for our optimal map and regression model

Our investigation showed that imputation map 28 along with *linear-priming*⊗*linear-PP-I* regression model generates the best predictive model for the NK cell fold expansion at day 9. Here we evaluate the errors in the regression coefficients for the above map and the model. The regression coefficients can be used to interpret relevant STAT-NF-κB synergy/ antagonism that regulates the weighted averaged NK cell fold expansion across donors (***Y*** in **Table S1B**). First, we chose all 12 treatment cytokine conditions (Fig. 1B) and applied our best predictive model and map (design matrix **X** of size 12×25) on weighted average (***Y***) (**Table S1B**) to evaluate the regression coefficients of vector ***β*** of STAT-STAT or STAT-NF-κB interactions between priming and PP-I period. Using L1 regularization, we selected the predictor variables whose coefficients were non-zero. We found 21 predictor variables are important (non-zero values) among 25 and discarded the predictor variables whose coefficients were zero in the design matrix **X**.

We computed the fluctuations in the **β** coefficients using a bootstrapping method, where we statistically generated *n* (=1000) number of NK cell fold expansion ***Y_Bootstrap_***≡***Ỹ*** vectors, each of which includes 12 conditions, by bootstrapping technique. To do that, we randomly selected N_A_(=10) donors from the donor 1 (D1) to donor 10 (D10) for treatment conditions 1-6 (**Table S1A**). Then, we calculated variances (σ_i_^2^) in NK cell fold expansion across conditions 1-6 for each of these N_A_ donors and calculated weight ω_i_ (i=1,…,N_A_) for each of them based on their deviances (deviance_i_), as 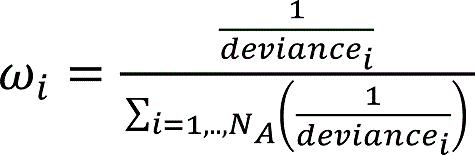(46), Where *deviance* of i^th^ donor, σ_i_ = (σ^2^_i_ − *medme*)^2^, where *medme*= [0.1 × *mean*(<σ_1_^2^, σ_2_^2^, …, σ_NA_^2^ =) + 0.9 × *median*(<σ_1_^2^, σ_2_^2^, …, σ_NA_^2^ =)]. The weighted average of fold expansion for *α*^th^ cytokine condition is calculated as Ŷ represents the fold expansion for *i*^th^ donor treated with *α*^th^ cytokine condition. Thus, we generated each bootstrapped, where *f* represents the fold expansion for *α*^th^ for *i*^th^ donor from **Table S1A**. Following the above method, we created ***Ỹ_B_*** (of size 2 × 1) for cytokine cocktail treatment conditions 7 and 8 by selecting N_B_(=4) donors randomly from donor 11 to donor 14. Similarly, we generated ***Ỹ_C_*** (of size 4 x 1) for cytokine cocktail treatment conditions 9-12 and by selecting N_C_(=3) donors randomly from donor 15 to donor 17. Next, we combined ***Ỹ_A_, Ỹ_B_*** and ***Ỹ_C_*** to create a new vector ***Ỹ*** (of size 12× 1) for all observed cytokine cocktail conditions. We use **X** (of size 12×21) and bootrapped weighted average ***Ỹ*** to calculate ***β*** (of size 21×1). We repeat this for *n* times, to get *n* number of ***β_j_*** (j=1,2,…,n). Then evaluated the ratio of mean (<***β***>) and standard deviation (***σ_β_***) for each element of {***β_j_***} and interpret synergy and antagonism for those predictor variables where (<β>/σ_β_) > 2 (26).

#### Leave out one Cross-validation for calculating prediction error for model selection

Given the NK proliferation fold expansion for 12 cytokine cocktail treatments, we find the overall cross-validation error by dividing the data into training data of size 11 (that includes 11 cytokine conditions) and test data of size 1 (the excluded 1 cytokine condition) (Fig. 1B). We use multiple linear regression (with LASSO or, Ridge regularizations) (26) to predict the NK proliferation fold expansion *Ŷ_α_* for α^th^ cytokine cocktail treatment condition. The total prediction error is calculated by summing over all 12 test sets, each representing an observed condition (Fig. 1B): Residual Sum of Squares (RSS) =∑ _α=1,…,12_ (*Y_α_-Ŷ_α_*)^2^. To predict the optimal cytokine condition for typical responders, we selected the imputation map with the minimum RSS (Fig. 3D**, 3E, Table S3** and **Fig. S1**).

#### Calculating R-square as a measure of overall prediction ability across 12 cytokine cocktail conditions

We have weighted average of fold expansion (***Y***) across donors, for 12 conditions (see Fig. 1B and **Table S1A**). By performing LOOCV, we predicted weighted fold expansion (*Ŷ_α_*) for each condition α. Where, α=1, 2,…,12. Then for each combination of imputation map and regression model, we calculated R^2^ between true (***Y***) and predicted values (***Ŷ***) of weighted average of fold expansion from all 12 conditions by obtaining Pearson Correlation coefficient (ρ), such that R^2^=ρ^2^. Comparisons between the above models for LOOCV in terms of RSS are shown in **Fig. S1** and **Table S3**.

## Data and code availability

All the codes are written in MATLAB software. Codes describing our in-silico models are available at the <u>link</u> https://github.com/indraniny?tab=repositories

## Author Contributions

SIK and JD planned the research. JD and IN developed computational method. IN processed the data and developed the in silico predictive framework. IN and RB made the figures. RB designed the *in vitro* experiments, collected, and analyzed the experimental data. RB, IN, SIK and JD wrote the manuscript.

## Inclusion and Diversity

We support inclusive, diverse, and equitable conduct of research.

## Acknowledgement

This work was supported by the NIH awards R01-AI 146581 to JD CRUK Cancer Research UK funding through the HUNTER HCC expediter network award (C9380/A18084) (SK), Miltenyi Biotech and by the Research Institute at the Nationwide Children’s Hospital. We thank Shashank Muniyappa for pointing us to RRelief. Thanks to Darren Wethington for discussions related to the model development. We acknowledge Ohio Supercomputer Center for usage of MATLAB.

