## Supplemental Materials for "Impact of IL-21 on Natural Killer cell proliferation and function – a mathematical and functional assessment"

### Supplementary Text 1. RRelief correctly chooses the ground truth model

We applied RRelief to short-list imputation maps that are consistent with the data. Here we performed a sanity check for the RRelief algorithm [27], where we generated synthetic data from a ground truth model and evaluated if RRelief is able to select the ground truth model.

We created a ground truth model of weighted average of NK cell fold expansion $Y_{GT} = X\hat{\beta}$ where the regression coefficients $\{\hat{\beta}_j\}$ are estimated using the design matrix X for the optimal map (map 28) and the model (*linear-priming* ⊗ *linear-PP-I* model) and the weighted average (Y) of fold expansion data (of size 12 × 1) (see Table S1B). $Y_{GT}$ provides synthetic data for the weighted average of the fold expansion for 12 treatment conditions. Next, we applied RRelief analysis on $Y_{GT}$ to evaluate the weights for each of STATs/NF-κB for priming and PP-I period varying the design matrix X for 64 imputation maps (see Table S2). This was carried out with constraints that were used in our map selection process with the experimental data (see Fig. 3A in the main text). The constraints are the following: (i) Cytokine cocktail treatment condition 2 activates the STAT3 and (ii) For NK cell expansion, the activation of STAT3 and NF-κB plays a positive role (considered positive scores in RRelief).

- By varying 64 imputation maps, we calculated 64 weight vectors from $Y_{GT}$=MW, where M is a matrix of size 12 x 10. Each row of M represents a cytokine cocktail condition. First and last 5 columns represent the presence of STAT1, STAT3, STAT4, STAT5 and NF-κB in the priming and in the PP-I period, respectively. The elements of matrix M changes with the imputation maps (see Table S2).
- Each weight vector W (of size 10 × 1) gives a score [-1,+1] related to each feature variables (STAT and NF-κB). We calculated the weights for each imputation map varying the nearest neighbor number (k).

Finally, we selected the imputation maps which show positive scores for STAT3 and NF-κB in both priming and PP-I period by varying nearest neighbor number (k=2,3,…,10). This filtering method provided map 28 (ground truth model) along with other imputation maps 8, 14, 16, 18, 32, 42, 44, 58 and 60. This provides a sanity check on RRelief method and indicates that our best predictive model (X) constructed with imputation map 28 has captured the effect of STAT3 and NF-κB inhibitors on fold expansion (see Fig. 3A in main text).

The code is available at GitHub:
https://github.com/indraniny/RRelief_ground_truth_Map28_L1_11Aug_23

**Supplementary Figures**

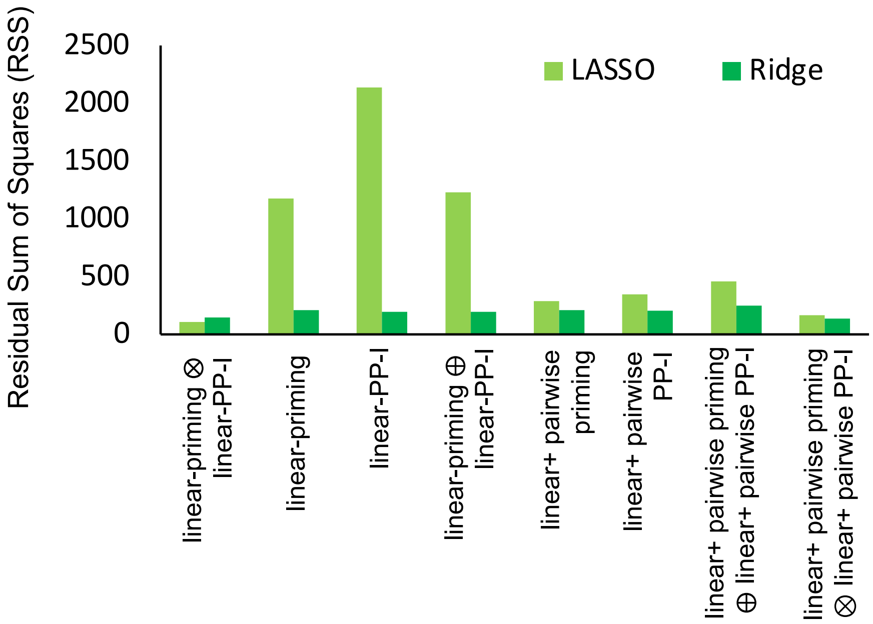


**Figure S1, related to Figure 3D. Comparison of prediction errors to choose the optimal map and regression model.** RSS (Residual Sum of Squares) as a measure of prediction error in weighted average of NK fold expansion (see ***Y*** in **Table S1B, Table S3**) for various regularized regression models. Height of each bar represents RSS across 12 LOOCV test sets, each representing a cytokine cocktail condition (**Fig. 1B** in main text). Light and dark green colors represent the regression model with LASSO (L1) and Ridge (L2) regularization, respectively (26). The left most regression model *(linear-priming* ⊗ *linear-PP-I)* with L1 regularization (light green) gives the minimum RSS, or the best prediction of fold expansion.

**RSS shown here, for each regularized regression model, are estimated for the optimal imputation map (varying maps 8, 14, 16, 28, 42, 44 to find the minimum RSS, See **Fig. 3D** in main text). For Ridge or Lasso regularization, the regularization constant λ is varied (λ=1e-11,1e-6,1e-3,…,1.0) to choose an optimal λ that gives minimum RSS.

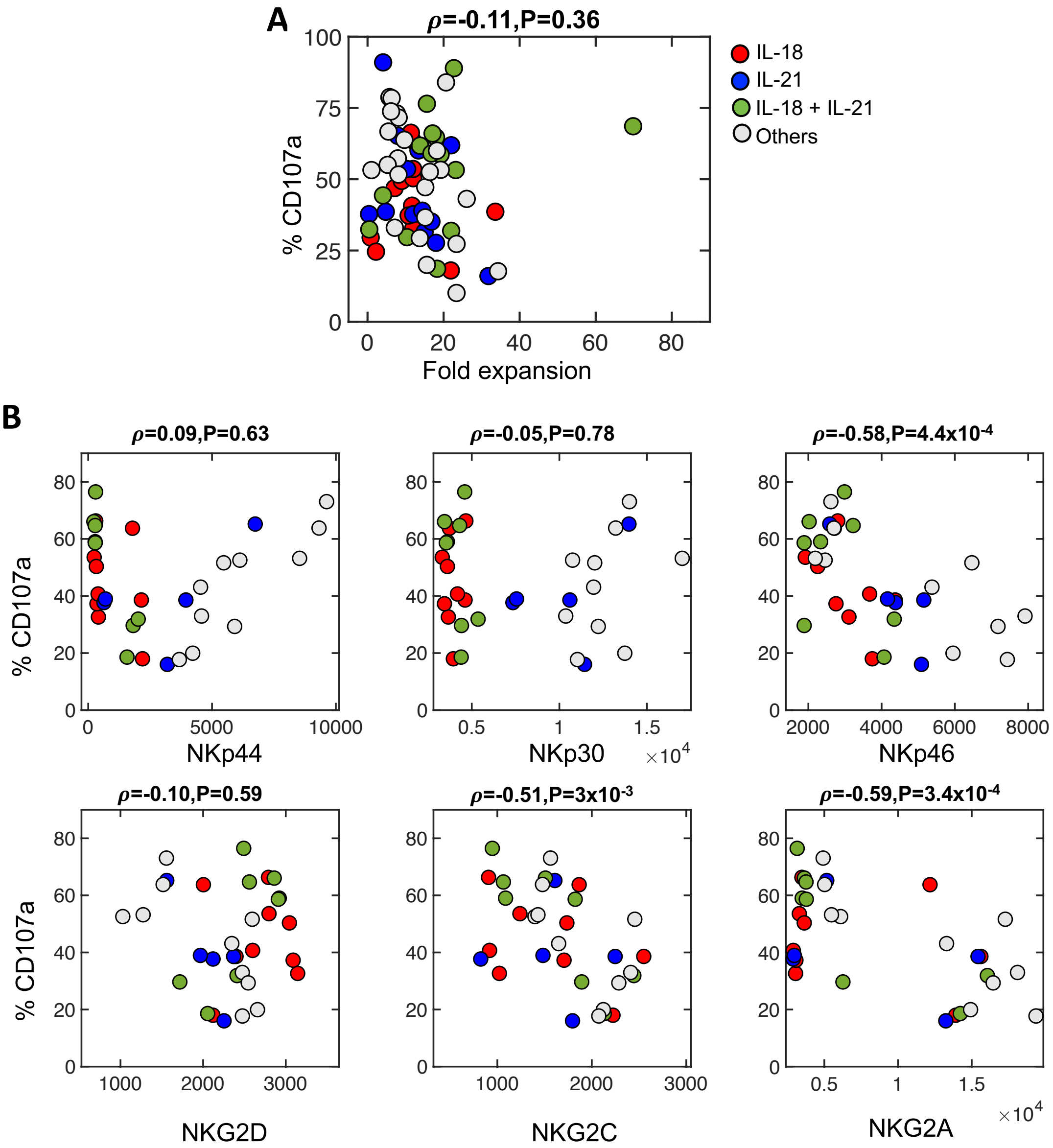


**Figure S2, related to Figure 5. Effect of IL-18 and IL-21 on NK cell fold expansion, cytotoxicity, and cell receptor expressions against hepatocellular carcinoma cell line HepG2**. All data shown here are for high CD16 expressing donors. High and low CD16 are gated based on the median of CD16 mean expressions from 11 donors (D3-D10, D15-D17) treated with 10 cytokine conditions (conditions 1-6,9-12). **(A)** Percentage of CD107a at day 10 is plotted against the NK fold expansion at day 9. Data are labelled based on the treatment conditions in the presence of either IL-18 (red) or, IL-21 (blue) or, both IL-18+21 (green) or none (grey). **(B)** Percentage of CD107a expressions is plotted against absolute cell receptor expressions. Pearson correlations (ρ) and *P* values are shown between %CD107a and each of absolute receptor expressions. NKp46, NKG2C and NKG2C show negative correlation with %CD107a.

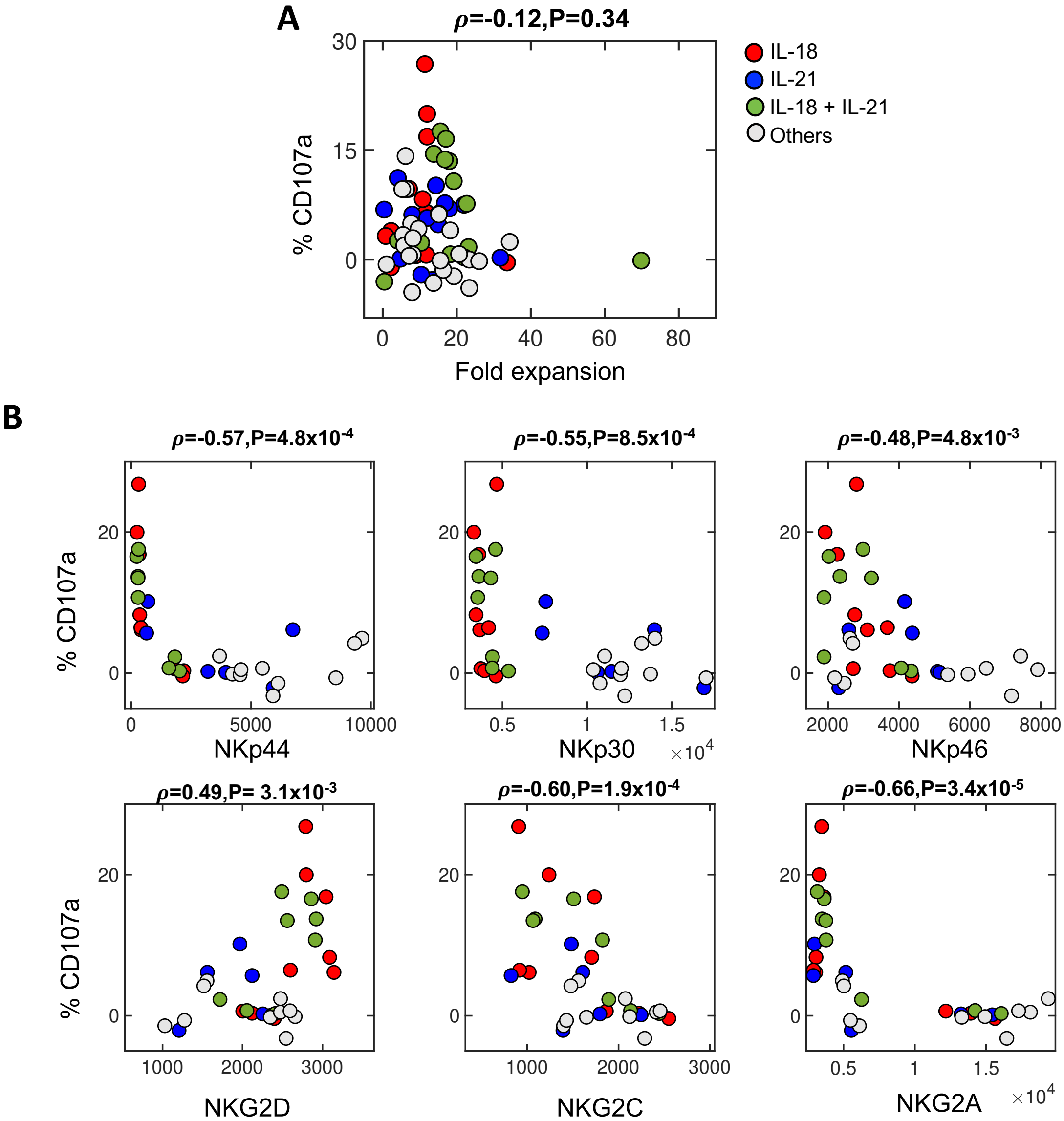


**Figure S3, related to figure 5. Effect of IL-18 and IL-21 on NK cell fold expansion, cytotoxicity, and cell receptor expressions against hepatocellular carcinoma cell line PLC**. All data shown here are for high CD16 expressing donors. High and low CD16 are gated based on the median of CD16 mean expressions from 11 donors (D3-D10, D15-D17) treated with 10 cytokine conditions (conditions 1-6,9-12). **(A)** Percentage of CD107a at day 10 is plotted against the NK fold expansion at day 9. Data are labelled based on the treatment conditions in the presence of either IL-18 (red) or, IL-21 (blue) or, both IL-18+21 (green) or none (grey). **(B)** Percentage of CD107a expressions is plotted against absolute cell receptor expressions. Pearson correlations (ρ) and *P* values are shown between %CD107a and each of absolute receptor expressions. All receptor expressions show negative correlation with %CD107a.

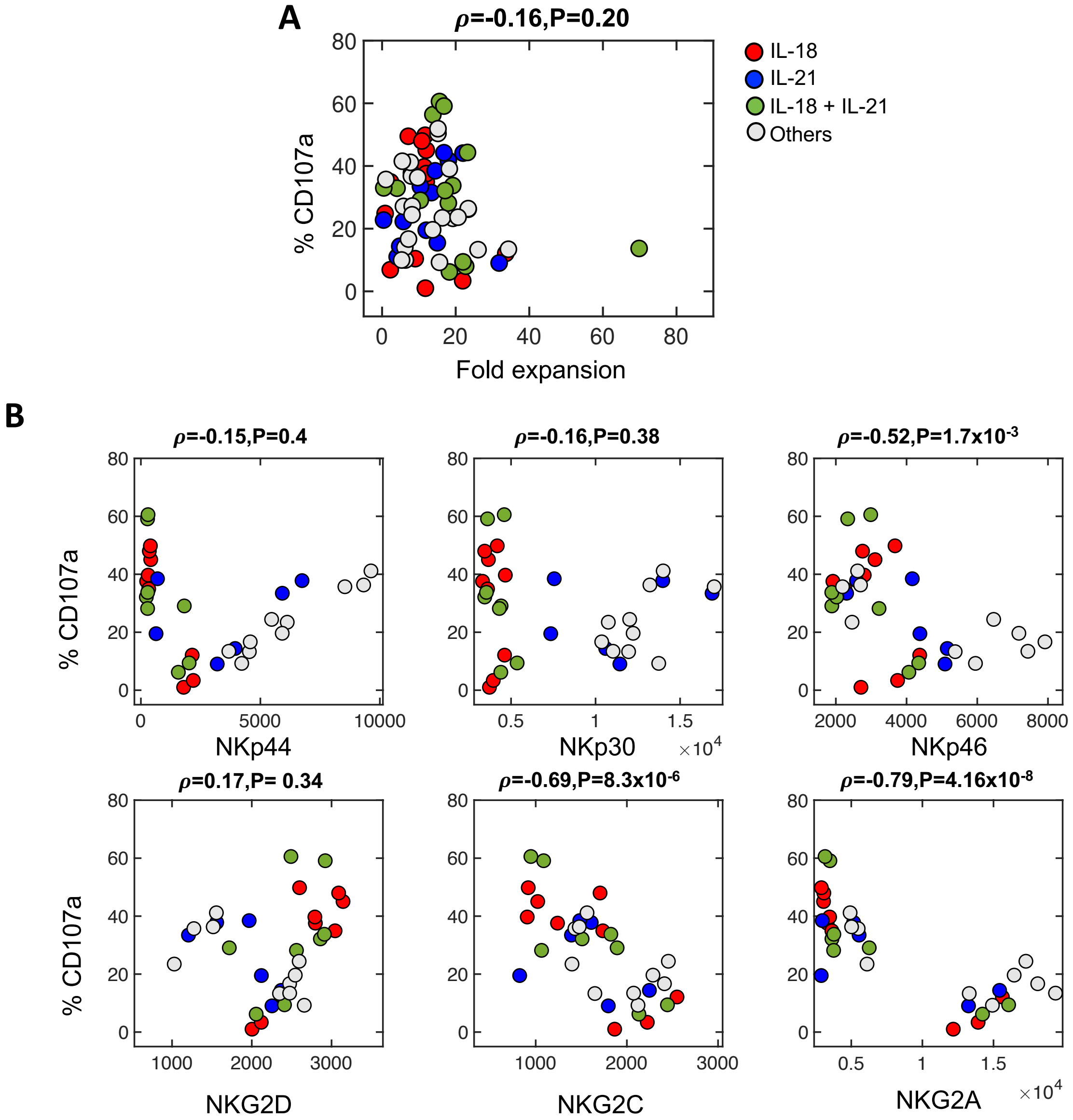


**Figure S4, related to figure 5. Effect of IL-18 and IL-21 on NK cell fold expansion, cytotoxicity and cell receptor expressions hepatocellular carcinoma cell line SNU475**. All data shown here are for high CD16 expressing donors. High and low CD16 are gated based on the median of CD16 mean expressions from 11 donors (D3-D10, D15-D17) treated with 10 cytokine conditions (conditions 1-6,9-12). **(A)** Percentage of CD107a at day 10 is plotted against the NK fold expansion at day 9. Data are labelled based on the treatment conditions in the presence of either IL-18 (red) or, IL-21 (blue) or, both IL-18+21 (green) or none (grey). **(B)** Percentage of CD107a expressions is plotted against absolute cell receptor expressions. Pearson correlations (ρ) and *P* values are shown between %CD107a and each of absolute receptor expressions. NKp46, NKG2C and NKG2C expressions show negative correlation with %CD107a.

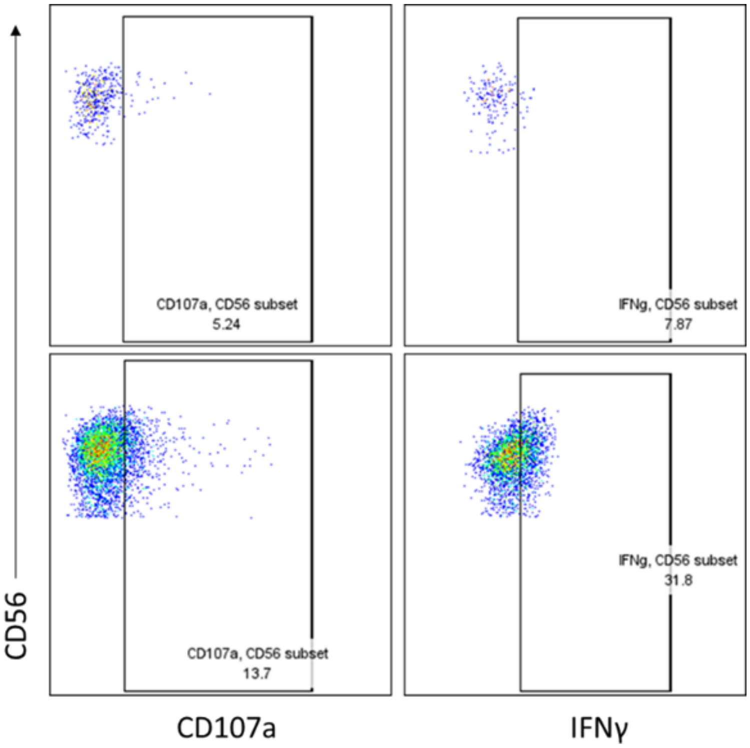


**Figure S5. Gating strategy to identify CD107a$^+$ NK during degranulation, using flow cytometry.**

NK cells were isolated from PBMC and cultured with different cytokines combinations for 10 days. NK cells were incubated without target cells (NT= no target) to detect the baseline degranulation of NK cells. The same gating was applied when NK cells were incubated with target cells (HepG2, as shown by the representative plot. The same gating was applied to the other cell lines used as targets, PLC, SNU475, Huh7). The final CD107a$^+$ NK cells with each target cell line were calculated by subtracting from it the NT CD107a$^+$ NK. The same gating strategy was applied to all the donors used.

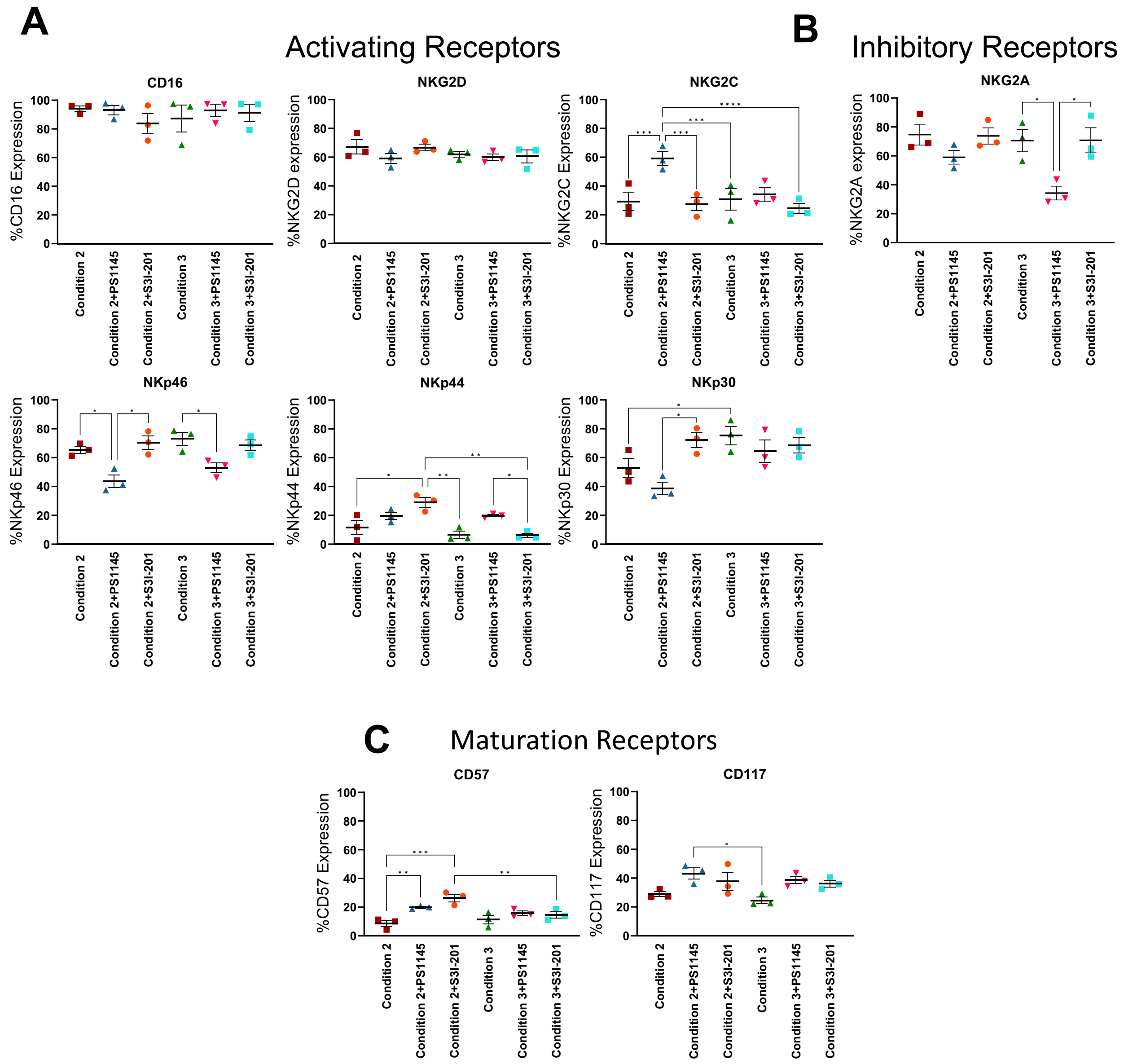


**Figure S6, related to figure 4. Effect of STAT3 and NF-κB inhibition on the phenotype of NK cells**

**(A)** Activating, **(B)** inhibitory, **(C)** maturation receptors expressed at day 10 of *in vitro* culture by cytokine-activated NK cells, cultured in the presence of STAT3 and NF-κB inhibitor during the 16hr priming stage. Data shown as mean of 3 different donors ± SEM. Data analyzed by RM one-way ANOVA, and Tukey multiple comparison correction. (*P≤0.05, ***P*≤0.01, ****P*≤0.001, *****P*≤0.0001).

**Supplementary Tables**

**NK cell fold expansion data at day 9 used to inform model:**

**Table S1A: NK cell fold expansion across donors for various cytokine cocktail treatment conditions**

| Treatment \ Donor | D1 | D2 | D3 | D4 | D5 | D6 | D7 | D8 | D9 | D10 |
|---|---|---|---|---|---|---|---|---|---|---|
| Condition 1 | 1.45 | 3.15 | 23.4 | 23.4 | 7.96 | 7.7 | 5.8 | 6.34 | 26.07 | 13.74 |
| Condition 2 | 1.75 | 1.29 | 2.28 | 7.12 | 0.82 | 2.22 | 9 | 11.76 | 21.9 | 33.6 |
| Condition 3 | 2.5 | 3.63 | 4.06 | 23.2 | 0.46 | 10.38 | 69.84 | 22.71 | 18.3 | 21.99 |
| Condition 4 | 0.95 | 2.24 | 13.4 | 22 | 10.42 | 7.98 | 5.76 | 4.1 | 31.8 | 4.8 |
| Condition 5 | 1.24 | 2.65 | 19.24 | 15.18 | 1.04 | 9.68 | 8.18 | 6.18 | 15.6 | 8.13 |
| Condition 6 | 1.35 | 2.32 | 18.26 | 15.18 | 16.38 | 5.5 | 20.64 | 5.3 | 34.32 | 7.17 |

| Treatment \ Donor | D11 | D12 | D13 | D14 |
|---|---|---|---|---|
| Condition 7 | 10.2 | 12 | 4.68 | 9 |
| Condition 8 | 9.6 | 15 | 6 | 11.1 |

| Treatment \ Donor | D15 | D16 | D17 |
|---|---|---|---|
| Condition 9 | 15 | 12 | 0.4 |
| Condition 10 | 18 | 14.4 | 16.8 |
| Condition 11 | 12 | 11.7 | 10.8 |
| Condition 12 | 13.8 | 15.6 | 16.8 |

**Table S1B: Weighted and standard average of NK cell fold expansion across donors for various cytokine cocktail conditions**

| | Weighted average ($\boldsymbol{Y}$) | Standard average ($\bar{\boldsymbol{Y}}$) |
|---|---|---|
| Condition 1 | 11.94 | 11.90 |
| Condition 2 | 13.63 | 9.17 |
| Condition 3 | 20.55 | 17.71 |
| Condition 4 | 11.85 | 10.34 |
| Condition 5 | 8.68 | 8.71 |
| Condition 6 | 13.35 | 12.64 |
| Condition 7 | 7.31 | 8.97 |
| Condition 8 | 8.80 | 10.42 |
| Condition 9 | 14.61 | 9.13 |
| Condition 10 | 17.55 | 16.40 |
| Condition 11 | 11.96 | 11.50 |
| Condition 12 | 14.03 | 15.40 |

**Table S2: Possible imputation Maps or induced STATs/NF-κB transcription factors by Interleukins used in our in silico model**

| Imputation Map Index | IL-2 | IL-12 | IL-15 | IL-18 | IL-21 |
|---|---|---|---|---|---|
| 1 | STAT5 | STAT4 | STAT5 | NF-κB | STAT3 |
| 2 | STAT5 | STAT4 | STAT5 | NF-κB, STAT3 | STAT3 |
| 3 | STAT5 | STAT4 | STAT5 | NF-κB | STAT3, STAT1 |
| 4 | STAT5 | STAT4 | STAT5 | NF-κB, STAT3 | STAT3, STAT1 |
| 5 | STAT5, STAT1 | STAT4 | STAT5 | NF-κB | STAT3 |
| 6 | STAT5, STAT1 | STAT4 | STAT5 | NF-κB, STAT3 | STAT3 |
| 7 | STAT5, STAT1 | STAT4 | STAT5 | NF-κB | STAT3, STAT1 |
| 8 | STAT5, STAT1 | STAT4 | STAT5 | NF-κB, STAT3 | STAT3, STAT1 |
| 9 | STAT5, STAT3 | STAT4 | STAT5 | NF-κB | STAT3 |
| 10 | STAT5, STAT3 | STAT4 | STAT5 | NF-κB, STAT3 | STAT3 |
| 11 | STAT5, STAT3 | STAT4 | STAT5 | NF-κB | STAT3, STAT1 |
| 12 | STAT5, STAT3 | STAT4 | STAT5 | NF-κB, STAT3 | STAT3, STAT1 |
| 13 | STAT5, STAT4 | STAT4 | STAT5 | NF-κB | STAT3 |
| 14 | STAT5, STAT4 | STAT4 | STAT5 | NF-κB, STAT3 | STAT3 |
| 15 | STAT5, STAT4 | STAT4 | STAT5 | NF-κB | STAT3, STAT1 |
| 16 | STAT5, STAT4 | STAT4 | STAT5 | NF-κB, STAT3 | STAT3, STAT1 |
| 17 | STAT5, NF-κB | STAT4 | STAT5 | NF-κB | STAT3 |
| 18 | STAT5, NF-κB | STAT4 | STAT5 | NF-κB, STAT3 | STAT3 |
| 19 | STAT5, NF-κB | STAT4 | STAT5 | NF-κB | STAT3, STAT1 |
| 20 | STAT5, NF-κB | STAT4 | STAT5 | NF-κB, STAT3 | STAT3, STAT1 |
| 21 | STAT5, STAT3, STAT1 | STAT4 | STAT5 | NF-κB | STAT3 |
| 22 | STAT5, STAT3, STAT1 | STAT4 | STAT5 | NF-κB, STAT3 | STAT3, |
| 23 | STAT5, STAT3, STAT1 | STAT4 | STAT5 | NF-κB | STAT3, STAT1 |
| 24 | STAT5, STAT3, STAT1 | STAT4 | STAT5 | NF-κB, STAT3 | STAT3, STAT1 |
| 25 | STAT5, STAT4, STAT1 | STAT4 | STAT5 | NF-κB | STAT3 |
| 26 | STAT5, STAT4, STAT1 | STAT4 | STAT5 | NF-κB, STAT3 | STAT3 |
| 27 | STAT5, STAT4, STAT1 | STAT4 | STAT5 | NF-κB | STAT3, STAT1 |
| 28 | STAT5, STAT4, STAT1 | STAT4 | STAT5 | NF-κB, STAT3 | STAT3, STAT1 |

| | | | | | |
|---|---|---|---|---|---|
| 29 | STAT5, NF-κB, STAT1 | STAT4 | STAT5 | NF-κB | STAT3 |
| 30 | STAT5, NF-κB, STAT1 | STAT4 | STAT5 | NF-κB, STAT3 | STAT3 |
| 31 | STAT5, NF-κB, STAT1 | STAT4 | STAT5 | NF-κB | STAT3, STAT1 |
| 32 | STAT5, NF-κB, STAT1 | STAT4 | STAT5 | NF-κB, STAT3 | STAT3, STAT1 |
| 33 | STAT5, STAT4, STAT3 | STAT4 | STAT5 | NF-κB | STAT3 |
| 34 | STAT5, STAT4, STAT3 | STAT4 | STAT5 | NF-κB, STAT3 | STAT3 |
| 35 | STAT5, STAT4, STAT3 | STAT4 | STAT5 | NF-κB | STAT3, STAT1 |
| 36 | STAT5, STAT4, STAT3 | STAT4 | STAT5 | NF-κB, STAT3 | STAT3, STAT1 |
| 37 | STAT5, STAT3, NF-κB | STAT4 | STAT5 | NF-κB | STAT3 |
| 38 | STAT5, STAT3, NF-κB | STAT4 | STAT5 | NF-κB, STAT3 | STAT3 |
| 39 | STAT5, STAT3, NF-κB | STAT4 | STAT5 | NF-κB | STAT3, STAT1 |
| 40 | STAT5, STAT3, NF-κB | STAT4 | STAT5 | NF-κB, STAT3 | STAT3, STAT1 |
| 41 | STAT5, STAT4, NF-κB | STAT4 | STAT5 | NF-κB | STAT3 |
| 42 | STAT5, STAT4, NF-κB | STAT4 | STAT5 | NF-κB, STAT3 | STAT3 |
| 43 | STAT5, STAT4, NF-κB | STAT4 | STAT5 | NF-κB | STAT3, STAT1 |
| 44 | STAT5, STAT4, NF-κB | STAT4 | STAT5 | NF-κB, STAT3 | STAT3, STAT1 |
| 45 | STAT5, STAT4, STAT3, STAT1 | STAT4 | STAT5 | NF-κB | STAT3 |
| 46 | STAT5, STAT4, STAT3, STAT1 | STAT4 | STAT5 | NF-κB, STAT3 | STAT3 |
| 47 | STAT5, STAT4, STAT3, STAT1 | STAT4 | STAT5 | NF-κB | STAT3, STAT1 |
| 48 | STAT5, STAT4, STAT3, STAT1 | STAT4 | STAT5 | NF-κB, STAT3 | STAT3, STAT1 |
| 49 | STAT5, STAT3, STAT1, NF-κB | STAT4 | STAT5 | NF-κB | STAT3 |
| 50 | STAT5, STAT3, STAT1, NF-κB | STAT4 | STAT5 | NF-κB, STAT3 | STAT3 |
| 51 | STAT5, STAT3, STAT1, NF-κB | STAT4 | STAT5 | NF-κB | STAT3, STAT1 |
| 52 | STAT5, STAT3, STAT1, NF-κB | STAT4 | STAT5 | NF-κB, STAT3 | STAT3, STAT1 |

| | | | | | |
|---|---|---|---|---|---|
| 53 | STAT5, STAT4, STAT3, NF-κB | STAT4 | STAT5 | NF-κB | STAT3 |
| 54 | STAT5, STAT4, STAT3, NF-κB | STAT4 | STAT5 | NF-κB, STAT3 | STAT3 |
| 55 | STAT5, STAT4, STAT3, NF-κB | STAT4 | STAT5 | NF-κB | STAT3, STAT1 |
| 56 | STAT5, STAT4, STAT3, NF-κB | STAT4 | STAT5 | NF-κB, STAT3 | STAT3, STAT1 |
| 57 | STAT5, STAT4, STAT1, NF-κB | STAT4 | STAT5 | NF-κB | STAT3 |
| 58 | STAT5, STAT4, STAT1, NF-κB | STAT4 | STAT5 | NF-κB, STAT3 | STAT3 |
| 59 | STAT5, STAT4, STAT1, NF-κB | STAT4 | STAT5 | NF-κB | STAT3, STAT1 |
| 60 | STAT5, STAT4, STAT1, NF-κB | STAT4 | STAT5 | NF-κB, STAT3 | STAT3, STAT1 |
| 61 | STAT5, STAT4, STAT1, NF-κB, STAT3 | STAT4 | STAT5 | NF-κB | STAT3 |
| 62 | STAT5, STAT4, STAT1, NF-κB, STAT3 | STAT4 | STAT5 | NF-κB, STAT3 | STAT3 |
| 63 | STAT5, STAT4, STAT1, NF-κB, STAT3 | STAT4 | STAT5 | NF-κB | STAT3, STAT1 |
| 64 | STAT5, STAT4, STAT1, NF-κB, STAT3 | STAT4 | STAT5 | NF-κB, STAT3 | STAT3, STAT1 |

**Table S3: Comparison of RSS and overall fold expansion predictability of 12 cytokine cocktail conditions ($R^2$) between regression models**

**A: LASSO regularization**

| **Design matrix for Regression Model** | **Optimal Imputation Map** | **Number of predictors (p) in design matrix X** | **$R^2$** | **RSS** |
|---|---|---|---|---|
| *linear-priming* ⨂ *linear-PP-I* | 28 | 25 | 0.55 | 105 |
| *linear-priming* | 8 | 5 | 0.055 | 1176 |
| *linear-PP-I* | 8 | 5 | NAN | 2136 |
| *linear-priming* ⨁ *linear-PP-I* | 8 | 10 | 0.01 | 1230 |
| *linear+pairwise-priming* | 42 | 15 | 0.022 | 286.744 |
| *linear+pairwise-PP-I* | 42 | 15 | 0.075 | 346.97 |

| | | | | |
|---|---|---|---|---|
| *linear+pairwise-priming* ⊕ *linear+pairwise-PP-I* | 42 | 30 | 0.029 | 456.56 |
| *linear+pairwise-priming* ⊗ *linear+pairwise-PP-I* | 28 | 225 | 0.122 | 166.69 |

**B: Ridge regularization**

| **Design matrix for Regression Model** | **Optimal Imputation Map** | **Number of predictors (p) in design matrix X** | **$R^2$** | **RSS** |
|---|---|---|---|---|
| *linear-priming* ⊗ *linear-PP-I* | 8 | 25 | 0.179 | 146.709 |
| *linear-priming* | 42 | 5 | 0.029 | 209.97 |
| *linear-PP-I* | 8 | 5 | 0.077 | 194.59 |
| *linear-priming* ⊕ *linear-PP-I* | 8 | 10 | 0.001 | 194.93 |
| *linear+pairwise-priming* | 42 | 15 | 0.029 | 210.09 |
| *linear+pairwise-PP-I* | 8 | 15 | 0.02 | 202.94 |
| *linear+pairwise-priming* ⊕ *linear+pairwise-PP-I* | 8 | 30 | 0.001 | 249.73 |
| *linear+pairwise-priming* ⊗ *linear+pairwise-PP-I* | 8 | 225 | 0.026 | 138.009 |

**Table S4: STAT-STAT and STAT-NF-κB synergy between priming and PP-I**

| Variable in Linear regression model | $\langle\beta\rangle/\sigma_\beta$ |
|---|---|
| **$S_1\tilde{S}_1$** | **2.2396** |
| $S_1\tilde{S}_3$ | -0.2233 |
| $S_1\tilde{S}_4$ | -1.9139 |
| **$S_1\tilde{S}_5$** | **2.6298** |
| $S_1\tilde{S}_b$ | -1.4216 |
| **$S_3\tilde{S}_1$** | **6.042** |
| $S_3\tilde{S}_3$ | 0.0385 |
| **$S_3\tilde{S}_4$** | **-2.3203** |
| $S_3\tilde{S}_5$ | 1.5327 |
| **$S_3\tilde{S}_b$** | **2.4948** |
| $S_4\tilde{S}_1$ | -1.5219 |
| **$S_4\tilde{S}_3$** | **-4.2445** |
| **$S_4\tilde{S}_4$** | **4.2445** |
| $S_4\tilde{S}_b$ | 1.3767 |
| $S_5\tilde{S}_3$ | 0.3846 |
| $S_5\tilde{S}_4$ | 1.0845 |

| | |
|---|---|
| $S_5\tilde{S}_b$ | 0.3916 |
| **$S_b\tilde{S}_1$** | **-2.4783** |
| $S_b\tilde{S}_3$ | 1.2028 |
| $S_b\tilde{S}_4$ | -1.2028 |
| $S_b\tilde{S}_b$ | 1.2028 |

** $S_i$ and $\tilde{S}_i$ represent induced STAT/NF-κB during the priming or PP-I period, respectively.
IL-21 in priming results in the activation of STAT3 and STAT1 in the priming and which contributes to the term $\mathbf{S_1\tilde{S}_1}$ , $\mathbf{S_1\tilde{S}_5}$ , $\mathbf{S_3\tilde{S}_1}$ , $\mathbf{S_3\tilde{S}_4}$ , $\mathbf{S_3\tilde{S}_b}$ significantly and synergistically (positive value) except the $\mathbf{S_3\tilde{S}_4}$ term (negative value). This quantification of synergy or antagonism might not reflect the true STAT or NF-κB interactions that regulate NK cell fold expansions.

**Table S5: Correlation between %CD107a and NK receptor expressions aganist HCC cell lines**

| Cell lines | Receptors with positive Pearson correlation (ρ) with %CD107a | Receptors with negative Pearson correlation (ρ) with %CD107a | Correlation between NK cell fold expansion (day 9) and %CD107a expressions |
|---|---|---|---|
| Huh7 | NKp44, NKp30 | NKG2D, NKG2C, NKG2A | ρ= - 0.29, P = 0.04 |
| HepG2 | | NKp46, NKG2C, NKG2A | ρ= - 0.11, P = 0.36 |
| PLC | NKG2D | NKp44, NKp30, NKp46, NKG2C, NKG2A | ρ= - 0.12, P = 0.34 |
| SNU475 | | NKp46, NKG2C, NKG2A | ρ= - 0.16, P = 0.20 |

** For the receptors in 2$^{nd}$ and 3$^{rd}$ column, Pearson Correlation (ρ) are associated with P values < 0.05.
